# RNAPII and XPC remodel the 3D genome for UV repair

**DOI:** 10.64898/2026.06.23.734090

**Authors:** Veysel Oğulcan Kaya, Mustafa Malkoç, Lucian-Alexandru Todirică, Ogün Adebali, Hanspeter Naegeli, Michelle N. Yancoskie

## Abstract

The three-dimensional (3D) genome architecture is highly elastic, adapting to nuclear processes such as transcription and the DNA damage response^1,2^. Nucleotide excision repair (NER) acts within this chromatin context to detect and repair mutagenic lesions induced by ultraviolet (UV) irradiation^3^. UV irradiation has been shown to induce restructuring of the 3D genome across multiple scales, including chromatin compartments, domains, and loops. However, the extent to which NER activity contributes to this remodelling remains unresolved, as the only prior study tracking such UV-induced changes was limited to repair-proficient cells^4^. Here, by combining genome-wide chromatin profiling of repair-deficient human cells with loop extrusion simulations, we show that the transcription-coupled NER initiator RNA polymerase II (RNAPII) and the global-genome NER initiator Xeroderma pigmentosum group C (XPC) each constrain loop extrusion upon UV irradiation. These events counter the loop-lengthening effects of UV-induced transcriptional shutdown, leading to shorter chromatin loops and reinforced chromatin domains that facilitate efficient lesion recognition and repair. The contribution by RNAPII extends its role beyond activating transcription-coupled repair to promoting a genome-wide repair-permissive state. The contribution by XPC establishes 3D genome reorganisation as an active mechanism both initiated and harnessed by DNA repair, rather than a passive consequence of DNA damage. Together, these findings advance our understanding of how nuclear processes coordinate on and alter a shared chromatin substrate to preserve genome integrity.

## Introduction

The three-dimensional (3D) genome is continuously remodelled during routine nuclear processes such as transcription, replication, recombination, and repair^1,2^. Our understanding of genome reorganisation during the DNA damage response stems largely from studies of DNA double-strand breaks (DSBs), which can be artificially introduced at defined genomic locations. Early studies of 3D genome restructuring during DSB induction reported increased insulation of self-interacting chromatin domains, termed topologically associating domains (TADs), following damage^5,6^. This reinforcement of pre-existing TADs was subsequently observed across additional cell types and DSB-inducing agents and was associated with enhanced repair, with DSB repair foci spreading from break sites until pre-existing TAD boundaries^5–8^. More globally, damaged TADs within active (A-compartment) chromatin were found to cluster together, which was postulated to concentrate repair factors and thereby increase their efficiency^8^.

In contrast to DSB induction, much less is known about how exposure to ultraviolet (UV) irradiation, a pervasive environmental mutagen, reshapes the 3D genome. Whereas engineered DSB systems permit chromatin changes to be tracked at reproducible damage sites^9^, no comparable strategies exist for UV-induced DNA lesions, which arise stochastically across the genome. UV irradiation induces helix-distorting photoproducts, primarily cyclobutane pyrimidine dimers (CPDs), resolved by the nucleotide excision repair (NER) pathway. Global-genome NER (GG-NER) processes the majority of these lesions through damage detection by DNA damage-binding protein 2 (DDB2) and Xeroderma pigmentosum group C (XPC),, whereas transcription-coupled repair (TC-NER) resolves the subset that stall elongating RNA polymerase II (RNAPII) within actively transcribed regions^3,10,11^.

Most work on UV-induced chromatin remodelling has focused on nucleosome-scale changes, with the ‘access-repair-restore’ model describing nucleosome repositioning and histone mark rewriting^12,13^. To date, we have published the only study tracking UV-induced large-scale changes. There, we observed early (12 min to 1 h post-UV) 3D genome reorganisation across multiple scales, including stronger segregation of chromatin into A and B compartments, increased insulation of pre-existing TADs, and strengthening of chromatin loops. The reinforcement of TADs and loops was associated with enhanced lesion repair, consistent with the notion that these architectural changes contribute functionally to the repair process rather than arising as a secondary consequence of UV exposure^4^.

The potential repair advantage conferred by genome reorganisation led us to consider whether cells actively promote these architectural changes, potentially through the repair machinery itself. In the present study, we characterised NER-associated contributions to genome restructuring by analysing the pre- and post-UV 3D genome of cells proficient and deficient for the GG-NER damage sensor XPC and the general NER factor Xeroderma pigmentosum group A (XPA). Before UV irradiation, XPC-deficient cells exhibited an unexpected decoupling between CCCTC-binding factor (CTCF) occupancy and chromatin insulation, which are typically positively associated^14–16^. This observation led us to investigate whether factors other than CTCF contribute to the architectural changes accompanying UV-induced DNA repair. Integrating chromatin profiles from repair-deficient cell lines with loop extrusion simulations revealed that lesion-stalled RNA polymerase II (RNAPII) and repair-associated barriers constrained loop extrusion in repair-proficient cells, counteracting the loop-lengthening effects of UV-induced transcriptional shutdown. Further constraining loop extrusion, the redistribution of XPC from TAD boundaries to UV-induced lesions was consistent with increased cohesin unloading as well as enhanced CTCF-mediated retention of cohesin at the boundaries.

Accordingly, the TADs showing the greatest loss of XPC and the greatest inferred loss of RNAPII after UV irradiation underwent the strongest reinforcement and, subsequently, the most efficient CPD repair. Thus, NER-associated processes promote a repair-permissive chromatin architecture characterised by reinforced TADs and shorter loops. Together, these findings support a model in which UV-induced genome restructuring constitutes an active component of the DNA damage response driven by the coupling of RNAPII stalling and repair initiation at lesions, rather than a passive reflex to DNA damage.

## Results

### UV irradiation triggers genome remodelling

To investigate how UV-induced DNA damage and subsequent repair activity alter chromatin architecture, we performed chromatin profiling of NER-proficient and NER-deficient U2OS cells before and after UV irradiation. Our genome-wide datasets include measurements of DNA damage, repair factor occupancy, chromatin accessibility, histone modifications, and (from Hi-C, a high-throughput chromosome conformation capture approach) chromatin contacts. These datasets span multiple resolutions, ranging from kilobase-scale chromatin loops and TADs to megabase-scale chromatin compartments (Fig. 1a; Extended Data Fig. 1–3).

**Figure 1:**
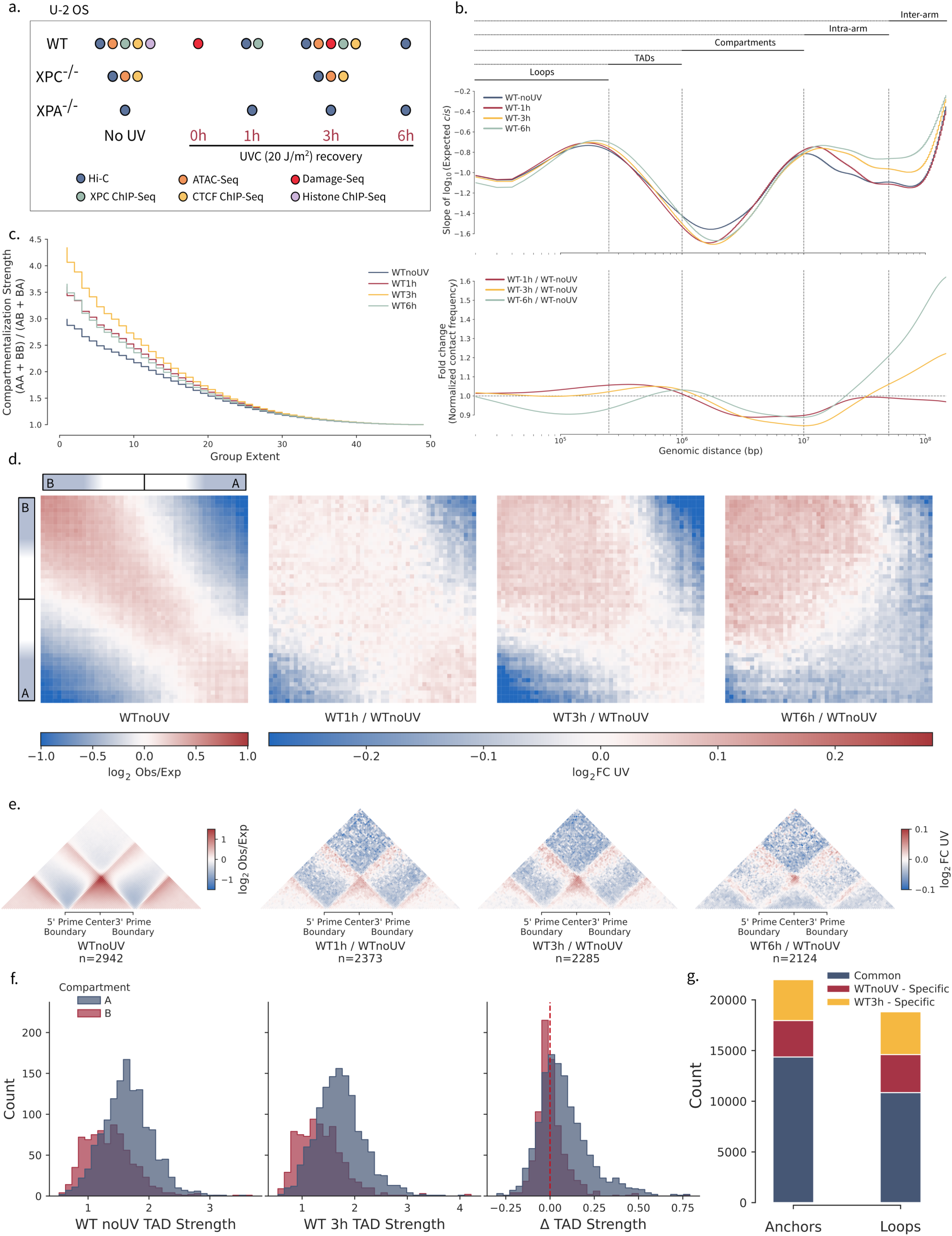
Experimental setup and genome reorganisation in wild-type cells. **a,** Experimental overview. Repair-proficient (wild-type, WT) and repair-deficient (*XPC*^-/-^, *XPA*^-/-^) U2OS cells were analysed before ultraviolet C (UVC) irradiation and at multiple post-UV (±20 J m^-2^) timepoints using Hi-C, genome-wide DNA damage mapping (damage-seq), chromatin profiling of XPC, chromatin accessibility (ATAC-seq), histone modifications (H3K27ac, H3K4me1, and H3K4me3), and CTCF occupancy. **b,** Genome-wide distance-decay profiles from WT cells of cis chromatin contacts derived from ICE-normalised Hi-C contact maps at 10-kb resolution, showing the slope of average contact probability, and its fold change relative to pre-UV, as a function of genomic distance. **c,** Compartment strength profiles in WT cells before and after UV irradiation calculated from cis contacts up to 100 Mb apart, showing enrichment of same-compartment over different-compartment contacts as a function of compartment rank (group extent), where lower extent corresponds to stronger compartment identity. **d,** Compartment interaction patterns in WT cells before and after UV irradiation, showing relative enrichment of same-compartment (A-A and B-B) versus inter-compartment (A-B) chromatin contacts for loci up to 100 Mb apart. **e,** Aggregate Hi-C contact maps centred on topologically associating domains (TADs) in WT cells, showing average chromatin interaction enrichment before UV, and the change after UV, across TAD bodies and flanking regions. Maps include flanking regions of one TAD length on each side. **f,** Distributions of TAD strength in WT cells before and 3 h after UV irradiation and the corresponding difference (3 h–pre-UV). TADs are grouped and coloured by compartment assignment. The dashed vertical line in the difference panel denotes no change (x = 0). **g,** Shared and timepoint-specific chromatin loops and loop anchors detected in WT cells before UV irradiation and 3 h post-UV.

From Hi-C on repair-proficient (wild-type; WT) cells, we observed a progressive increase in inter-chromosomal (trans) contacts following UV irradiation (Extended Data Fig. 1a), consistent with intermingling of chromosome territories upon genotoxic stress. Intra-chromosomal (cis) contact frequencies between loci separated by ∼10 kb–1 Mb became progressively enriched from 1–3 h post-UV, but were depleted by 6 h compared to pre-UV. By contrast, contacts spanning 1–10 Mb were depleted throughout the post-UV timecourse, and contacts spanning >10 Mb became enriched by 3–6 h (Fig. 1b). The transient enrichment in short- to mid-range contacts (10 kb–1 Mb) and concomitant reduction in long-range (1–10 Mb) contacts recapitulate trends previously observed in UV-irradiated HeLa cells, although those changes were only monitored up to 1 h post-UV^4^.

Contacts at genomic scales exceeding 1 Mb are associated with chromatin compartments, which are thought to be shaped largely by phase separation^17^. Compartment identity, whereby A-compartment is associated with active and B-compartment with inactive chromatin, was largely preserved across timepoints, with infrequent compartment switching (Extended Data Fig. 1d). This contrasts with the extensive B-to-A compartment transitions reported upon DSB induction, thought to facilitate repair by increasing accessibility to damaged loci^18^. Despite stable compartment identities, compartment-specific contacts were dynamically remodelled after UV exposure. For cis loci separated by up to 10 Mb, intra-compartment contacts (A-A and especially B-B) relative to inter-compartment (A-B) contacts increased up to 3 h post-UV (Fig. 1c,d; Extended Data Fig. 1e, 2a). Notably, enrichment of A-A contacts was most pronounced at early timepoints, whereas B-B contacts continued to increase at later timepoints. This temporal pattern mirrors the progression of UV lesion repair from active into inactive regions^19,20^. Accordingly, the sustained increase in B-B contacts at later timepoints may arise from DDB2-mediated decondensation of heterochromatin^21^ as repair progressively shifts towards inactive chromatin. At longer genomic distances spanning chromosome arms and chromosomes, A-A contacts decreased whereas B-B contacts, initially depleted, progressively increased. A-B contacts also increased, consistent with this shift in repair priority (Extended Data Fig. 2a).

Short- to mid-range contacts correspond to TADs and loops–structures generally attributed to loop-extrusion-dependent processes^22,23^. We quantified TAD strength, defined as the ratio of intra- to inter-TAD contacts^24^, across timepoints. Again consistent with the 3D genome changes captured in HeLa cells, TAD strength tended to increase after UV irradiation, especially within A-compartment regions (Fig. 1e,f; Extended Data Fig. 3a). Because TAD strengthening had plateaued by 6 h, we focused subsequent analyses on the 3 h timepoint as the window of maximal domain-level genome reorganisation. We found that most chromatin loops detected prior to UV exposure were maintained by 3 h post-UV, with loops gained after UV modestly outnumbering those that were lost (Fig. 1g). We next asked whether these changes corresponded to differential NER activity.

### Repair favours reinforced chromatin

To probe possible functional consequences of UV-induced chromatin reorganisation, we investigated whether the reinforcement of TADs observed after UV irradiation is associated with the formation and repair rates of cyclobutane pyrimidine dimers (CPDs), the predominant DNA lesions induced by UV irradiation. We mapped CPDs, captured by high-sensitivity damage-sequencing (damage-seq;^25^), across the TADs identified from Hi-C. Initial CPD formation at TAD boundaries and loop anchors was low, consistent with reported protection against CPD formation at CTCF-bound regions^26^. Because sequence-based simulation profiles also showed reduced vulnerability to CPDs at these sites, local DNA sequence composition contributed to the relatively reduced CPD levels (Extended Data Fig. 4a).

We stratified the TADs into quartiles according to their TAD strength 3 h after UV irradiation, with Q4 TADs becoming strongest–reflecting enhanced intra-TAD and reduced inter-TAD interactions–and Q1 becoming weakest (Extended Data Fig. 3c-e), and assigned loops to the quartile of their encompassing TAD. Consistent with HeLa cells, in U2OS cells the strongest pre-damage TADs further strengthened after UV irradiation, whereas the weakest further weakened (Extended Data Fig. 3e). This stratification was also associated with chromatin openness. The strongest TADs were enriched in A-compartment, aligning with their high pre-UV chromatin accessibility as measured by assay for transposase-accessible chromatin sequencing (ATAC-seq). The weakest TADs were associated with B-compartment and tended to be longer (Extended Data Fig. 3e). Following UV irradiation, the strongest TADs and their loops underwent the largest increase in CTCF occupancy, corresponding to their enhanced insulation (Fig. 2a,b). We found that the strongest TADs and especially their loops underwent enhanced CPD repair regardless of their initial damage burden (Fig. 2a, b). The prioritisation of early repair across the strongest TADs, which tend to lie in euchromatic regions, is consistent with preferential NER activity in open, transcriptionally active chromatin^19,20^.

**Figure 2:**
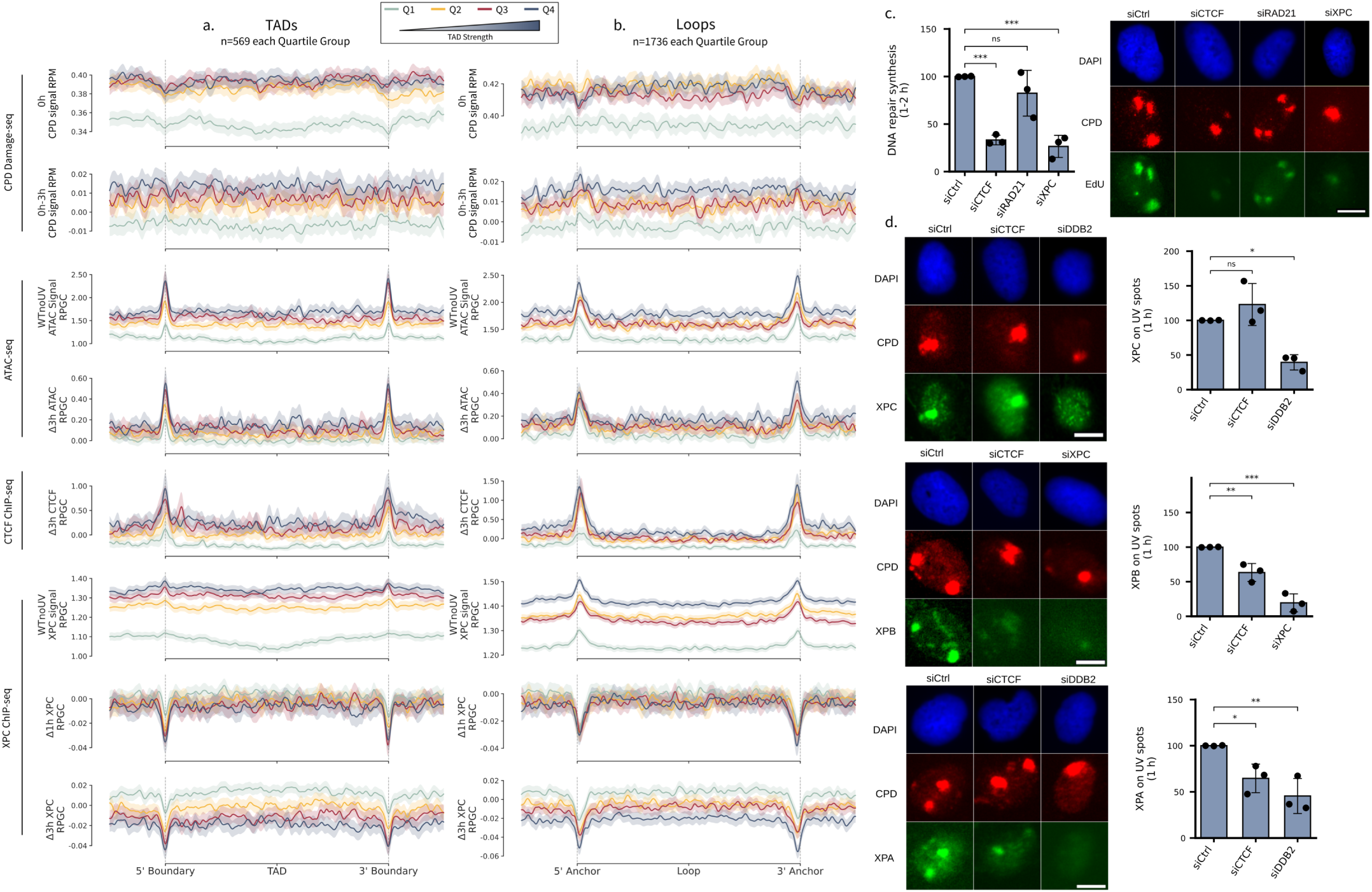
Strengthened TADs are associated with enhanced CPD repair after UV. **a,** Topologically associating domains (TADs) were grouped into quartiles based on 3 h post-UV TAD strength (Q4, strongest; Q1, weakest). Metaprofiles show cyclobutane pyrimidine dimer (CPD) damage-seq at 0 h and 0–3 h, chromatin accessibility (ATAC-seq) before and 3 h after UV, 3 h change in CTCF occupancy, and pre-damage, 1 h change, and 3 h change in XPC occupancy (ChIP-seq), across TADs and flanking regions. Dashed vertical lines denote TAD boundaries. **b,** Chromatin loops were assigned to the TADs in which they resided, downsampling to the loop count of the quartile containing the fewest loops. Metaprofiles show the same features as in a across loops for each group. Loop analyses are restricted to loops whose anchors fall within annotated TADs and for which ≥90% of the loop span is contained within a single TAD, entailing a subset of the loops depicted in Fig. 1. Dashed vertical lines denote loop anchors. Lines show mean signal across regions in panels a and b; shaded bands, mean ± SEM. **c,** DNA repair synthesis measured by EdU incorporation following local UV irradiation in cells treated with the indicated siRNAs. Representative immunofluorescence images (DAPI, CPD, EdU) are shown at the right; scale bars, 10 µm. Bars denote mean; error bars, SD; dots, independent biological experiments (n=3). Statistical significance was assessed with one-way ANOVA followed by a two-sided Dunnett’s multiple-comparison test comparing each experimental condition to the control. Significance: ns, P ≥ 0.05; ***P < 0.001. **d,** Quantification of XPC, XPB, and XPA accumulation at UV-induced DNA damage sites measured by immunofluorescence. Representative images (DAPI, CPD, and indicated repair factor) are shown at the right; scale bars, 10 µm. Bars denote mean, error bars, SD; dots, independent biological experiments (n=3). Statistical analysis was performed as in c. Significance: ns, P ≥ 0.05; *P < 0.05; **P < 0.01; ***P < 0.001.

Contrasting the enhanced CPD repair efficiency across stronger TADs and loops, the GG-NER factor XPC, an early sensor of CPD damage, preferentially redistributed to lower-quartile TADs within 1–3 h after UV irradiation. This XPC mobility away from pre-UV binding sites was especially prominent at TAD boundaries and chromatin loop anchors, where XPC is strongly enriched before damage (Fig. 2a,b; Extended Data Fig. 4b,c). Previous studies have likewise reported rapid XPC turnover and redistribution across chromatin following UV irradiation^25,27^. To test whether genome restructuring supports repair progression beyond XPC redistribution, we performed repair synthesis and immunofluorescence assays measuring EdU incorporation and repair factor occupancy, respectively, at CPD foci in UV-irradiated cells depleted by siRNA of architectural or repair factors. Acute CTCF depletion globally but reversibly abolishes TAD insulation and canonical (CTCF-cohesin) loops^15^. Acute depletion of the cohesin subunit RAD21 rapidly abolishes CTCF-anchored loops^28–30^. In our repair synthesis assay, CTCF depletion markedly inhibited repair within 2 h after UV exposure, comparable in magnitude to depletion of the core GG-NER factor XPC. In contrast, depletion of the cohesin subunit RAD21 had little effect on repair synthesis (Fig. 2c, Extended Data Fig. 5a-e). Through immunofluorescence assays, we detected XPC retention at CPD foci in CTCF-depleted cells. Depleting CTCF therefore inhibits NER without impairing initial damage sensing by XPC. Instead, recruitment of downstream NER factors XPB, a subunit of the TFIIH damage verification complex, and XPA, which stabilises the resulting single-stranded DNA complex, was impaired at 1–2 h post-UV (Fig. 2d; Extended Data Fig. 5f-h). The repair synthesis and XPC, XPB and XPA recruitment phenotypes of siCTCF cells match those observed upon depletion of the GG-NER factor and histone methyltransferase ASH1L, which facilitates repair progression by establishing a stable chromatin scaffold for NER factor binding^25,31^. Thus, chromatin destabilisation upon CTCF depletion is compatible with impaired retention of downstream repair machinery.

### XPC opposes chromatin insulation

We next asked whether the architectural rewiring observed after UV is mediated by the repair machinery itself, thus constituting an active component of the repair response rather than a mere byproduct of DNA damage. To this end, we performed Hi-C, CTCF chromatin immunoprecipitation sequencing (ChIP-seq), and ATAC-seq in *XPC*^-/-^ U2OS cells, which cannot initiate GG-NER and therefore cannot remove CPDs outside of the actively transcribed regions processed by transcription-coupled NER (TC-NER). We observed that *XPC*^-/-^ cells had more insulated chromatin than WT cells before damage and that, like WT cells, they responded to damage by becoming even more insulated. Notably, this inverse relationship between XPC abundance and 3D genome rigidity is echoed in pluripotent stem cells, which combine high XPC expression (with XPC transcriptionally coactivating OCT4 and SOX2^32–34^) and hypertranscription^35^ with a relaxed genome architecture, including weak TAD boundaries, compared to differentiated cells^36–38^.

*XPC*^-/-^ cells had higher baseline cis-compartment saddle strength than WT cells. Like WT cells, their long-distance (>100 Mb) B-B and A-B compartment contacts had increased by 3 h post-UV, with concomitant decrease in A-A (Extended Data Fig. 1e, 2b,d). Furthermore, *XPC*^-/-^ cells displayed features consistent with stronger loop extrusion both before and after UV irradiation. Before UV, they had more frequent short- to mid-range contacts, much stronger TADs, and more numerous and stronger chromatin loops than WT cells. By 3 h after UV, these short- to mid-range contacts had increased (Extended Data Fig. 1c), TADs and their boundaries had become stronger (Fig. 3a), and loop strength had increased modestly (Fig. 3b), with several thousand additional loops and loop anchors detected in *XPC*^-/-^ than in WT cells both before and after UV irradiation (Fig. 3c). By contrast, *XPA*^-/-^ cells, deficient for both GG-NER and TC-NER, tended to undergo compartment and domain strength weakening accompanied by an increase in trans contacts, consistent with secondary chromatin perturbations accompanying persistent unrepaired lesions (Extended Data Fig. 1, 2c-d, 3a).

**Figure 3:**
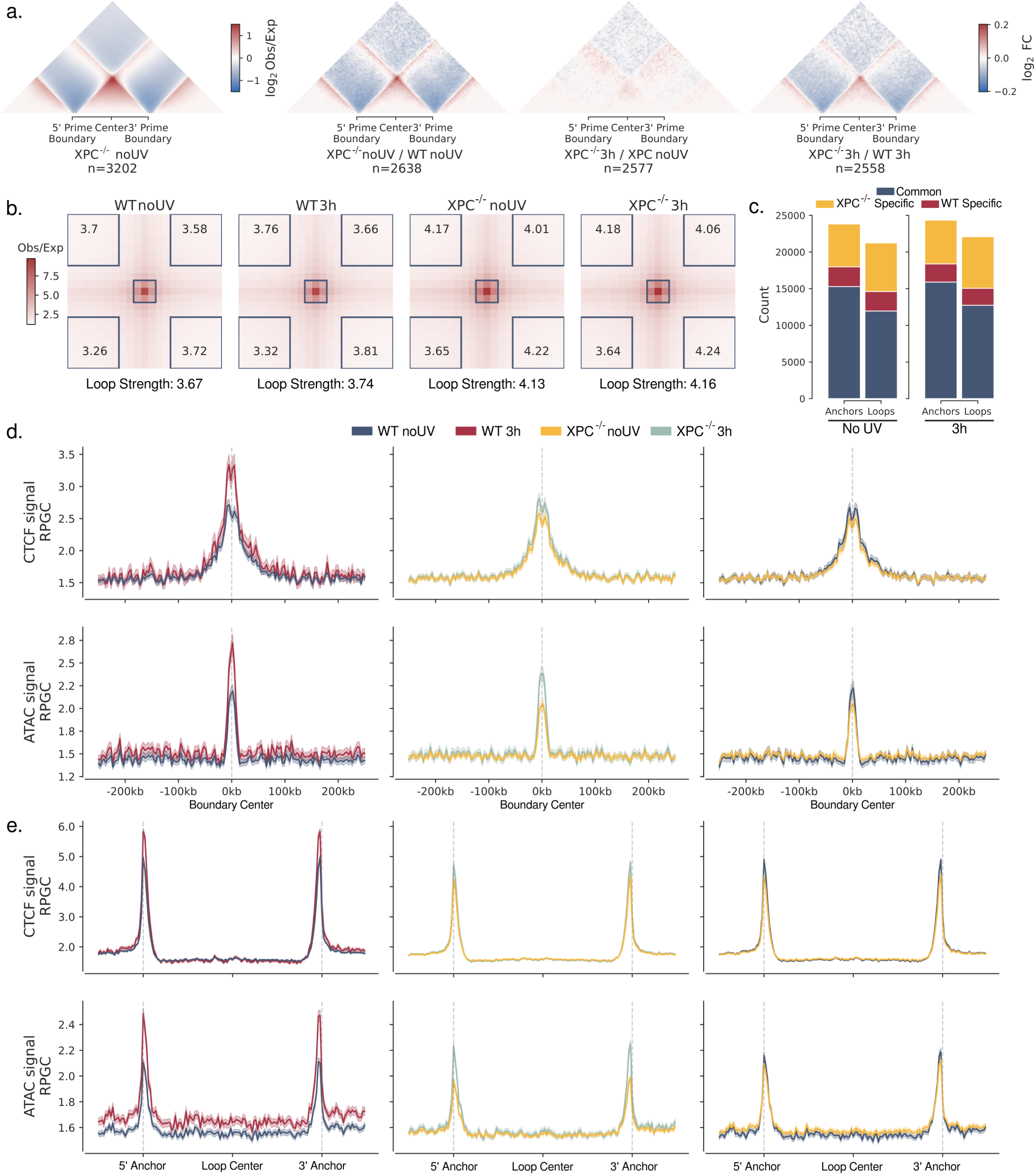
Altered chromatin organisation in XPC-deficient cells before and after UV irradiation. **a,** Aggregate Hi-C contact maps centred on topologically associating domains (TADs) in wild-type (WT) and *XPC*^-/-^ U2OS cells before and 3 h after UV irradiation. Difference maps show changes in contact frequency relative to untreated cells and between genotypes. Maps include flanking regions of one TAD length on each side. **b,** Aggregate Hi-C contact maps centred on chromatin loop anchors (loop pixels) in WT and *XPC*^-/-^ U2OS cells before and 3 h after UV irradiation. Corner values indicate central loop enrichment relative to the corresponding local background regions. Loop strength values below each map quantify central loop enrichment normalised to the average of the top left, top right, and bottom right background regions. **c,** Numbers of chromatin loops and loop anchors shared between WT and *XPC*^-/-^ cells or specific to each genotype before and 3 h after UV irradiation. **d,** Metaprofiles of CTCF ChIP-seq signal and chromatin accessibility (ATAC-seq) centred on TAD boundaries preserved between genotypes (left, middle columns) or timepoints (right column). UV irradiation enhances CTCF enrichment at maintained boundaries in both WT and *XPC*^-/-^ cells, whereas boundary insulation remains stronger in *XPC*^-/-^ cells despite comparable CTCF accumulation. **e,** Metaprofiles of CTCF ChIP-seq signal and chromatin accessibility (ATAC-seq) centred on chromatin loops preserved between genotypes or timepoints. Dashed vertical lines denote the 5′ and 3′ chromatin loop anchors. Lines show mean signal across regions in panels d and e; shaded bands, mean ± SEM.

CTCF occupancy modulates chromatin insulation in a dose-dependent manner. Convergently oriented CTCF at TAD boundaries and loop anchors poses a barrier to cohesin translocation, with tandem arrays of CTCF-bound sites bestowing stronger insulation^14–16,39^. We therefore examined whether altered CTCF binding could explain the stronger loop extrusion-associated features observed in *XPC*^-/-^ cells. CTCF ChIP-seq revealed an unexpected relationship between CTCF occupancy and insulation strength. Within each genotype, UV-induced strengthening of TADs was accompanied by increased CTCF enrichment, consistent with established models of CTCF-dependent insulation. In turn, the greater CTCF occupancy increase at TAD boundaries of WT cells is consistent with reports linking the NER endonucleases XPF and XPG to CTCF-associated looping^40,41^, as WT cells engage these factors more extensively than do TC-NER-restricted *XPC*^-/-^ cells. Moreover, boundaries and anchors within each cell type underwent a corresponding increase in chromatin accessibility, consistent with local UV-induced remodelling including nucleosome repositioning and altered protein occupancy^13,42^. Across genotypes, however, this relationship was inverted: at TAD boundaries preserved between untreated WT and *XPC*^-/-^ cells, *XPC*^-/-^ cells displayed lower CTCF occupancy and chromatin accessibility despite their stronger TADs (Fig. 3d). A similar pattern was observed across chromatin loops. Within each genotype, increased loop strength after UV coincided with increased CTCF enrichment and chromatin accessibility at anchors, yet across genotypes *XPC*^-/-^ cells had stronger loops despite less CTCF binding and lower chromatin accessibility (Fig. 3e). Thus, loop and TAD strengthening in *XPC*^-/-^ compared to WT cells cannot be explained solely by absolute local CTCF abundance, nor by chromatin accessibility.

Given the extensive co-localisation of XPC and CTCF both genome-wide (Extended Data Fig. 6a-b) and at TAD boundaries (Fig. 4a-b), significantly exceeding expectations from accessibility-matched, shu ed controls, we asked whether this surprising inverse relationship between CTCF abundance and insulation was also evident within untreated WT cells. We implemented a covariate-matched framework that stratified the genome into 10-kb bins containing at least one CTCF peak, and selected focal (high-XPC) and matched control (low-XPC) bins to maximise XPC contrast while controlling for chromatin accessibility and histone modification levels (H3K4me1/me3, H3K27ac from^25,43^) (Fig. 4c; Extended Data Fig. 6c-d). Despite lower CTCF occupancy, low-XPC regions had stronger insulation than focal regions (Fig. 4d-e), matching the *XPC*^-/-^ phenotype of stronger insulation despite reduced CTCF binding. Thus, partial absence of XPC (at select loci within XPC-proficient cells) is sufficient to recapitulate the apparent dampening of CTCF-mediated chromatin insulation.

**Figure 4:**
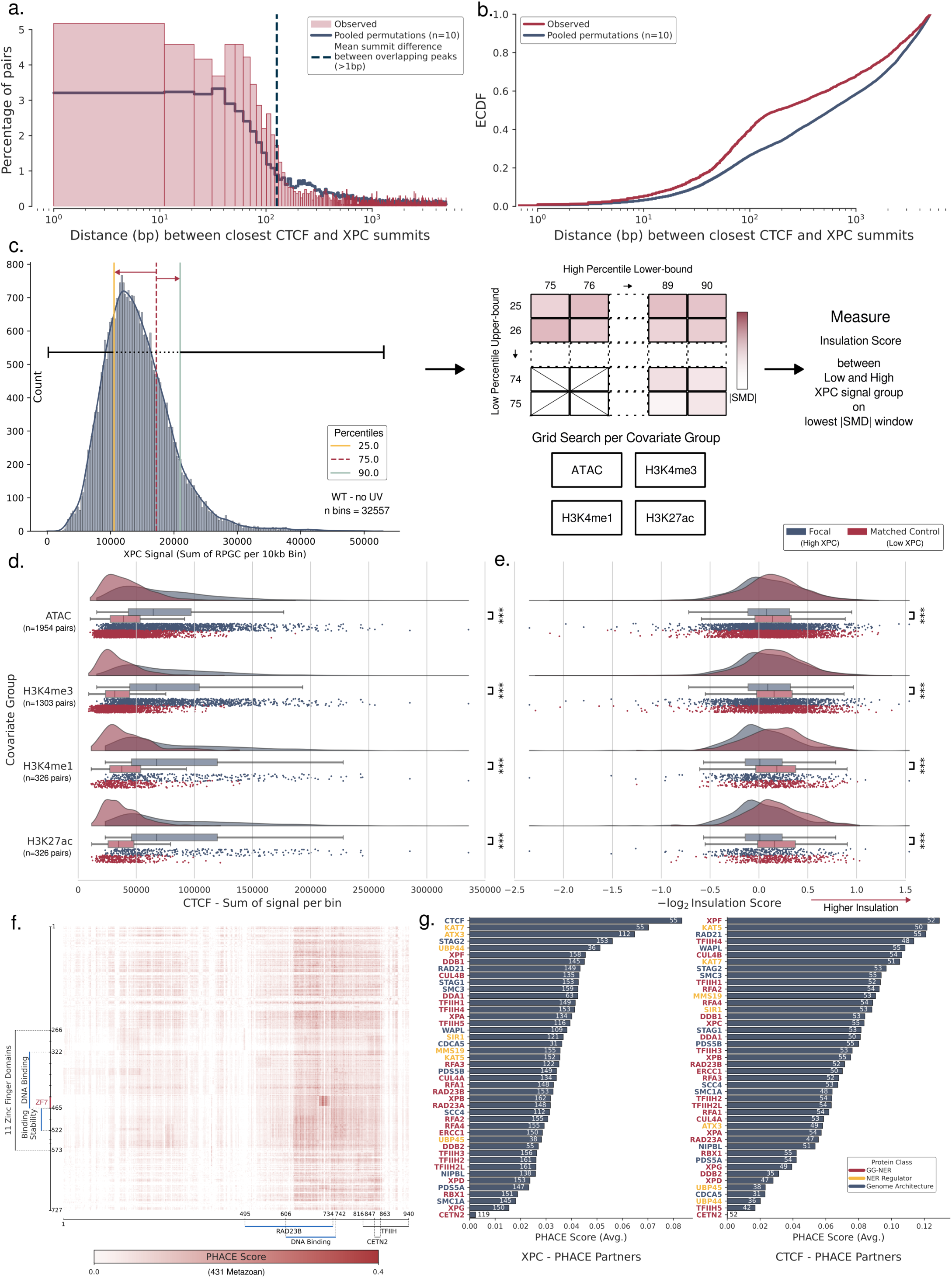
XPC occupancy is associated with reduced chromatin insulation in undamaged cells. **a,** Distribution of distances between the closest CTCF and XPC chromatin immunoprecipitation sequencing (ChIP-seq) peak summits at TAD boundaries in wild-type (WT) U2OS cells under untreated conditions. The observed distances (red hatched histogram) were compared to a null distribution (blue) generated by repeated permutations of XPC peak positions restricted to ATAC-seq-defined accessible chromatin while preserving chromatin identity, providing an accessibility-matched expectation for XPC-CTCF proximity. **b,** Empirical cumulative distribution function (ECDF) of observed (red) and permuted (blue) distances between CTCF and XPC peak summits. **c,** Schematic of the analytical framework used to define focal genomic regions with high XPC occupancy and matched control regions with low XPC occupancy. The genome was partitioned into 10-kb bins and stratified by XPC ChIP-seq signal (left), which was used to identify thresholds that maximise XPC contrast while maintaining comparable chromatin context. High- and low-XPC bins were then matched for chromatin covariates, including chromatin accessibility (ATAC-seq) and histone modification levels (H3K4me1/me3 and H3K27ac), defining focal and matched control sets for downstream analyses (right). **d,** Violin plots with overlaid boxplots show the distribution, median, and interquartile ranges of CTCF ChIP-seq signal across focal high-XPC and matched low-XPC control regions. Each data point denotes an individual 10-kb genomic bin, with n = 1,954, 1,303, 326, and 326 matched pairs for the ATAC, H3K4me3, H3K4me1, and H3K27ac covariate groups, respectively. Statistical significance was assessed by a two-sided Mann-Whitney U test; ***P < 0.001. **e,** Violin plots with overlaid boxplots show the distribution, median, and interquartile ranges of Hi-C insulation scores for the same focal and matched control regions, with the same sample sizes as in d. Statistical significance was assessed by a two-sided Mann-Whitney U test; ***P < 0.001. **f,** Residue-level phylogenetic analysis of co-evolution (with PHACE;^44^) heatmap between XPC and CTCF. Each pixel represents a pair of amino-acid positions between the two proteins. Signals in both proteins are concentrated at DNA-binding interfaces, including the RAD23B interaction interface in XPC and the seventh zinc finger of CTCF. **g,** Summary of top PHACE interaction partners for XPC (left) and CTCF (right) based on average PHACE scores across species. Bars are ranked by mean PHACE score, and numbers within bars indicate the number of eukaryotic species contributing to each estimate. The lower species counts for CTCF reflect its restriction to metazoans, whereas XPC orthologs are present across a broader set of eukaryotic species.

### XPC supports CTCF binding through TDG

To explore whether the inverse relationship between XPC presence and chromatin insulation reflects functional coupling of XPC with CTCF, we applied Phylogeny-Aware Detection of Molecular Coevolution (PHACE). PHACE tests whether amino acid substitutions in two proteins occur in a coordinated manner across independent evolutionary events^44^. Residue-level PHACE analysis identified discrete regions of evolutionary coupling between XPC and CTCF. Notably, the N-terminal YDF and KTYQR motifs of CTCF, despite their established role in modulating loop extrusion by stalling cohesin^39^, did not exhibit elevated PHACE signal relative to other domains of CTCF. Instead, the co-evolutionary signal in both proteins was concentrated at their DNA-binding domains, overlapping the RAD23B interaction interface of XPC^45^ and the seventh zinc finger (ZF7) of CTCF^46^ (Fig. 4f). ZF7 contacts CpG positions at the 5′ end of the CTCF motif. Methylation at this position strongly impairs CTCF binding. By contrast, methylation at CpG positions towards the 3′ end of the motif is better tolerated^47^. In parallel, XPC has been implicated together with RAD23B in facilitating thymine DNA glycosylase (TDG) turnover on chromatin, thereby accelerating completion of DNA demethylation^48^. Together, these observations are consistent with a role of XPC in facilitating methylation-sensitive CTCF binding by promoting efficient completion of TDG-dependent demethylation at CTCF motifs. This mechanism can explain the lower pre-damage CTCF ChIP-seq signal at TAD boundaries and loop anchors in *XPC*^-/-^ cells compared to WT cells (Fig. 3d-e), whereby the absence of XPC as a TDG displacer attenuates demethylation of the 5′ CpG within the CTCF motif, inhibiting CTCF binding.

We additionally examined PHACE signal between CTCF or XPC and other NER and chromatin architectural factors to assess whether the observed coupling extended across a broader network (Fig. 4g; Extended Data Fig. 6e). Summary PHACE analysis across NER factors, NER regulators, and chromatin architectural proteins identified CTCF as the highest-scoring co-evolution partner of XPC. The NER endonuclease XPF emerged as the top partner for CTCF, consistent with prior reports linking XPF to CTCF-associated chromatin looping^40,41^. Across the broader set of factors, CTCF exhibited co-evolutionary signals with multiple partners, consistent with its general role in chromatin regulation, but at residues distinct from those strongly coupled to XPC. Notably, XPC was among the partners with the strongest PHACE signal. Despite no direct XPC-CTCF interaction having been reported to date, these observations are consistent with shared evolutionary constraints linking XPC-associated DNA demethylation to methylation-sensitive CTCF binding.

### RNAPII overrides CTCF-mediated insulation

While XPC may promote CTCF binding through demethylation, this mechanism does not explain how XPC-deficient loci have stronger chromatin insulation despite reduced CTCF occupancy. We therefore considered whether the elevated insulation observed in *XPC*^-/-^ cells, and at XPC-deficient loci in untreated WT cells, arises through other 3D genome regulators. One possibility is that XPC influences the abundance or chromatin residence time of the loop extruder cohesin. Our PHACE analyses revealed strong co-evolutionary coupling between XPC and the cohesin subunits RAD21 and STAG2 (Fig. 4g). STAG1/2 and RAD21 form part of the cohesin interface that interacts with the unloading factor WAPL^49^. If XPC attenuates cohesin unloading, its absence could shorten cohesin residence time, limiting the distance over which cohesin extrudes and thereby biasing chromatin contacts towards shorter distances. This model is consistent with the enrichment of loops and short (≲500 kb) TADs that we observed in *XPC*^-/-^ compared to WT cells–especially for short, accessible TADs, as these were most enriched for XPC before damage (Extended Data Fig. 1c, 3e; Fig. 2a).

Another possibility is that XPC modulates the behaviour of RNA polymerase II (RNAPII), a semi-permeable barrier to cohesin. When stalled or terminated, RNAPII poses an additional obstacle to loop extrusion alongside the more static barrier posed by CTCF. During elongation, RNAPII asymmetrically modulates loop extrusion by one-sidedly promoting–or, depending on the direction of the encounter, impeding–cohesin translocation^50–53^. Through these interactions, elongating RNAPII can redistribute cohesin away from CTCF-bound sites and has been proposed to constitute the ‘kick’ that disrupts transient enhancer–promoter contacts in the ‘kick-and-kiss’ model of gene regulation^54^. Conversely, inhibition of RNAPII elongation promotes cohesin accumulation at CTCF-bound sites^50,51,53,55^. Such RNAPII-based mechanisms have been proposed to account for chromatin insulation patterns that are not readily explainable by local CTCF occupancy^51,53,56^.

Before UV irradiation, XPC functions as a cofactor of RNAPII at active promoters, where its depletion inhibits assembly of the preinitiation complex and attenuates transcription of a subset of genes^57^ (potentially contributing to the lower basal chromatin accessibility observed at TAD boundaries and loop anchors in *XPC*^-/-^ cells compared to WT (Fig. 3d–e)). Upon UV irradiation, there is widespread stalling of RNAPII at DNA lesions within gene bodies. The stalled RNAPII serves as the initiating signal for TC-NER and, to permit access to lesions by the repair machinery, must subsequently be cleared through backtracking, dissociation from chromatin, or proteasomal degradation^58,59^. Through a distinct pathway, promoter-bound RNAPII complexes, separate from lesion-stalled elongating RNAPII, are degraded within approximately 1 h of UV irradiation. This response transiently shuts down transcription, affording TC-NER time to remove lesions before new polymerases enter damaged gene bodies^58^.

Because XPC is an RNAPII cofactor, we asked whether altered RNAPII dynamics contributes to the 3D genome configuration of *XPC*^-/-^ cells, To this end, we re-analysed published Micro-C data from DLD-1 cells before and after degron-mediated RNAPII depletion^50^. We generated TAD boundary-centred maps and juxtaposed them with U2OS aggregate Hi-C maps displayed on the same scale. We identified two phenotypes that paralleled those of XPC-/- cells. First, RNAPII depletion reduced cross-TAD boundary contacts despite reduced CTCF binding (Fig. 5a,b). The TAD strengthening was greatest for TADs with the highest initial RNAPII occupancy (from^60^; Extended Data Fig. 7a–e). Second, building on the first phenotype, both RNAPII depletion and XPC deficiency increased stripe intensity (Extended Data Fig. 7b; Extended Data Fig. 10c). Stripe intensity reflects asymmetric loop extrusion, whereby cohesin is stalled by CTCF at only one TAD boundary such that the TAD is partially rather than fully extruded, a state that predominates *in vivo*^50,61–63^. Loss of elongating RNAPII, whether through depletion or (in *XPC*^-/-^ cells) through failure of the preinitiation complex to assemble, removes a moving to cohesin progression, allowing cohesin to reach TAD boundaries more readily. In turn, the reduced CTCF occupancy at these boundaries would favour one-sided stalling, thereby increasing stripe intensity.

**Figure 5:**
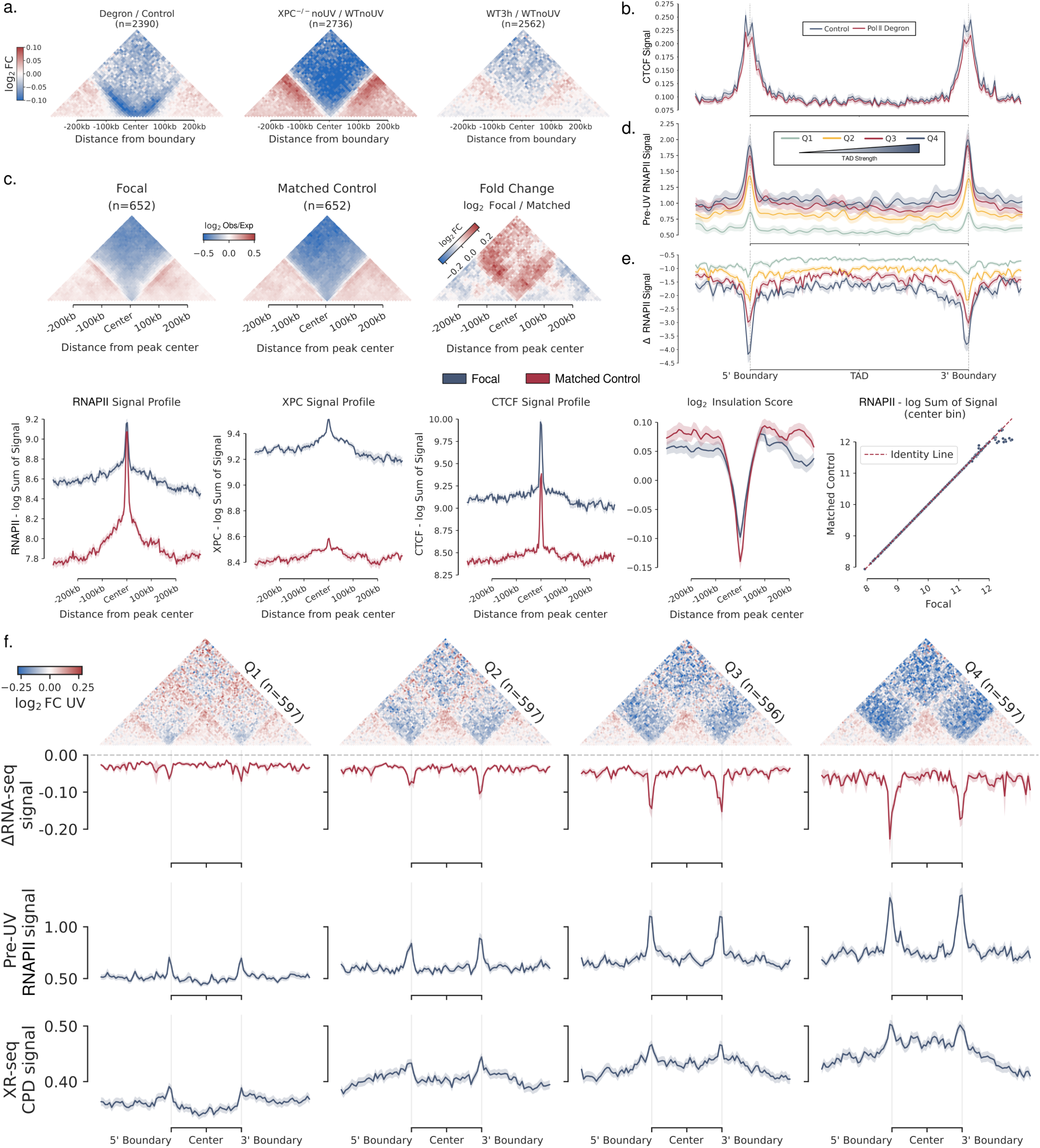
RNA polymerase II loss sharpens TAD insulation. **a,** Aggregate contact difference (fold change) maps centred on topologically associating domain (TAD) boundaries. Maps are shown for DLD-1 cells following degron-mediated RNA polymerase II (RNAPII) degradation (left; re-analysed from published degron Micro-C data;^50^), untreated *XPC*^-/-^ U2OS cells relative to untreated wild-type (WT) U2OS cells (centre), and WT U2OS cells 3 h after UV irradiation relative to untreated WT cells (right). **b,** Mean signal for CTCF^50^, centred on TADs preserved across control and RNAPII-degraded cells. **c,** Aggregate Hi-C contact maps centred on focal (high-XPC) and matched (low-XPC) 10-kb bins (averaged over a ±250-kb genomic interval) from the refined RNAPII-adjusted matched covariate set, corresponding difference map, and mean insulation, RNAPII, and XPC signal profiles (refined matching diagnostics in Extended Data Fig. 8). **d,** Pre-UV RNAPII occupancy (from^64^) across U2OS TADs stratified as in Fig. 2–that is, according to 3 h post-UV TAD strength (Q1–Q4, weakest to strongest). **e,** Change in RNAPII occupancy in lung-derived fibroblasts (MRC5) before and 75 min after UV irradiation (4 J m^-2^), projected onto TADs from a related fibroblast line (IMR90) stratified by TAD strength. **f,** TADs detected in HeLa cells^4^ were grouped into quartiles based on TAD strength 12 min after UV irradiation and profiled for changes in contact strength (12 min / pre-UV), transcriptional output (RNA-seq; first 12 min), pre-damage RNAPII occupancy, and repair of cyclobutane pyrimidine dimers (XR-seq; first 12 min). Line plots show mean signal across regions; shaded bands, mean ± SEM.

To test whether variation in RNAPII occupancy also contributes to insulation differences within non-challenged WT cells, we incorporated RNAPII occupancy from undamaged U2OS cells^64^ as an additional covariate into our matching framework. Although our covariate maps were not explicitly centred on TAD boundaries or loop anchors, XPC, CTCF, and RNAPII are all enriched at such elements together with active promoters. Accordingly, the regions flanking our central CTCF peak-containing bins provide a proxy for active TADs (Fig. 5c). Furthermore, because XPC, CTCF and RNAPII also co-localise genome-wide, focal (high-XPC) bins were intrinsically enriched for RNAPII relative to low-XPC controls. As this RNAPII enrichment extended beyond the central bin into the flanking regions, the focal group represents generally RNAPII-rich environments rather than a merely local difference at the central boundary-like element. Before matching for RNAPII, fold change (focal-over-control) aggregate Hi-C maps centred on the focal and matched 10-kb bins showed broad contact enrichment extending into both flanking regions surrounding (±250 kb) the central, CTCF peak-containing bin (Extended Data Fig. 8). After matching for central RNAPII occupancy, contacts spanning the central, boundary-like bin intensified (Fig. 5c). This pattern is consistent with multiple RNAPII molecules distributed throughout active TADs locally trapping cohesin after boundary bypass events, thereby limiting further extrusion and promoting short-range contacts across boundaries. Such contacts would weaken local insulation despite elevated CTCF occupancy.

### Repair creates transient barriers to extrusion

Disruption of productive RNAPII elongation accompanies both degron-mediated RNAPII depletion and UV irradiation, the latter causing both RNAPII stalling and degradation. Aggregate TAD boundary-centred Hi-C maps in WT U2OS cells 3 h after UV relative to untreated WT cells showed, like RNAPII-depleted cells, loss of cross-boundary contacts (Fig. 5a). To assess whether this association between RNAPII loss and domain strengthening extended beyond local TAD boundaries, we examined RNAPII distribution across the TAD quartiles from Fig. 2. While standard ChIP-seq cannot distinguish between different functional states of RNAPII, such as stalling versus elongation^65^, published pre-UV RNAPII ChIP-seq data from unirradiated U2OS cells^64^ showed the highest RNAPII occupancy across Q4 TADs and the lowest across Q1 (Fig. 5d). Although matching post-UV RNAPII data were unavailable to directly assess changes in RNAPII occupancy after damage, we projected RNAPII pre- and post-UV ChIP-seq data from MRC5 cells^58^ onto TAD strength-grouped quartiles from other lung-derived fibroblasts (non-challenged IMR90;^66^) and observed that Q4 TADs lost the most RNAPII signal (Fig. 5e). Furthermore, Q4 TADs in UV-exposed HeLa cells^4^ (defined by their 12-min post-UV TAD strength) had, as in U2OS, the highest cumulative pre-damage RNAPII load. Within 12 min of UV exposure, these Q4 TADs underwent the strongest reduction in transcriptional output as measured by RNA sequencing (RNA-seq). Like U2OS Q4 TADs, they were subject to the fastest repair, as measured by XR-seq^67^ (from^68^) (Fig. 5f).

Although RNAPII depletion and UV irradiation both reduced inter-TAD contacts, only UV irradiation enhanced short-range contacts. RNAPII depletion was accompanied by a loss of short-range (<500 kb) contacts (Fig. 6a). This trend is consistent with the longer loops reported in the RNAPII degron study and with the RNAPII barrier-removal model introduced above, in which RNAPII loss permits longer-range cohesin translocation^50,55^. We hypothesised that UV-induced stalling of RNAPII at lesions, which would introduce additional barriers to cohesin rather than remove them, accounts for the enrichment of short-range contacts observed after UV irradiation. Because the response to UV irradiation entails both RNAPII stalling and degradation, and because datasets capable of resolving the relative contributions of these processes were unavailable, we next turned to loop extrusion simulations. We modelled the main perturbations that accompany UV irradiation–RNAPII degradation (transcriptional arrest), TAD boundary strengthening, and lesion-associated barriers to cohesin extrusion–and examined their predicted effects on 3D genome organisation (Fig. 6b; Supplementary Methods for stochastic modelling of loop extrusion, transcription, and nucleotide excision repair). Given the limited experimental knowledge of several kinetic parameters expected to influence 3D genome organisation after UV irradiation, these simulations were intended to evaluate qualitative trends and mechanistic plausibility rather than to reproduce the exact magnitude of chromatin changes.

**Figure 6:**
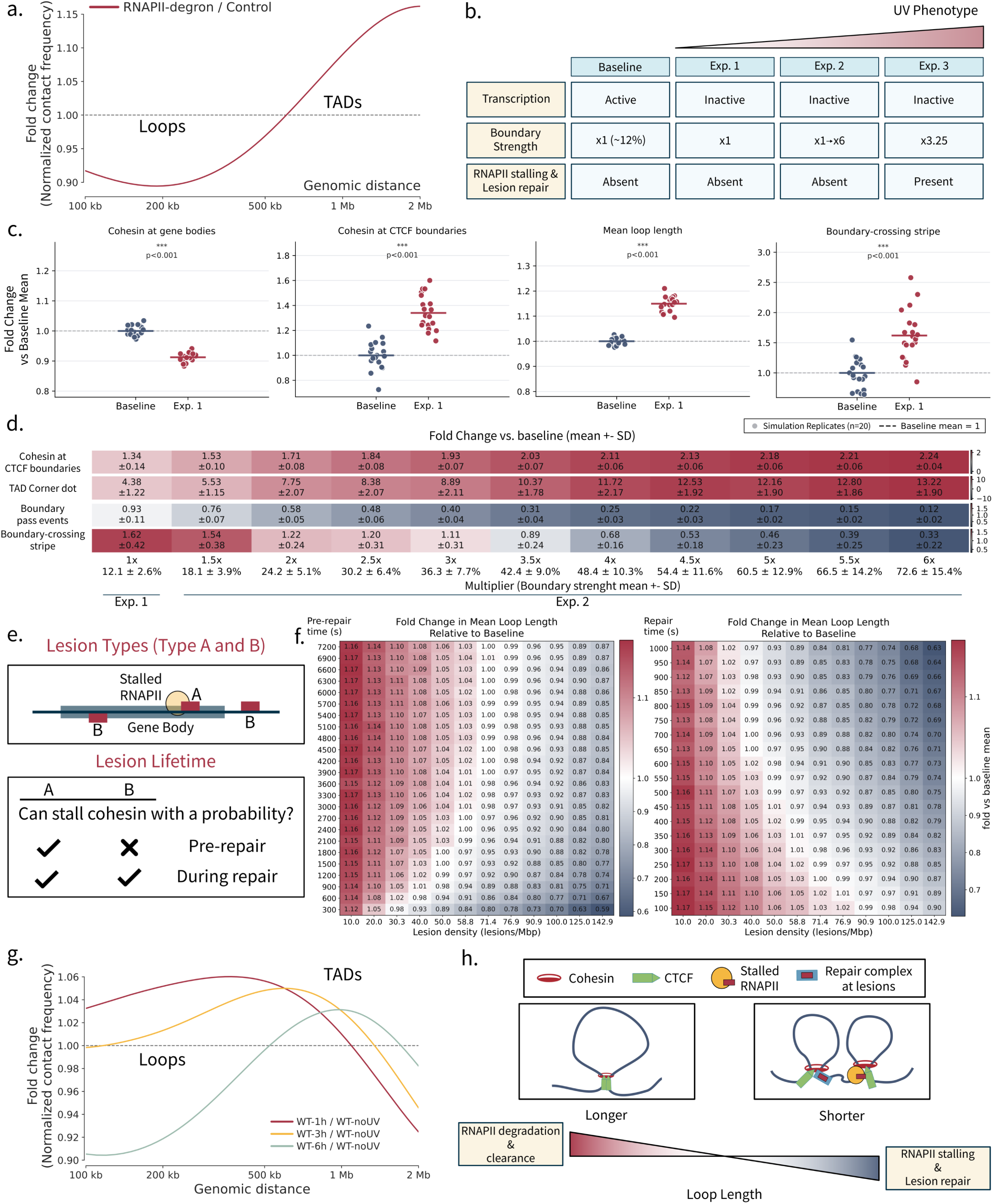
Loop extrusion simulations identify drivers of UV-induced genome restructuring. **a,** Genome-wide distance-decay profile from Micro-C of DLD-1 cells following degron-mediated RNA polymerase II (RNAPII) depletion (from^50^). Plots show the fold change in contact probability relative to control cells as a function of genomic distance, restricted to genomic separations of up to 2 Mb. **b,** Schematic of simulation conditions used to model progressive features of the UV response. The baseline condition represents unirradiated cells with active transcription, standard boundary strength (∼12% probability of cohesin crossing inhibition at CTCF boundaries, as in^61^), and no DNA lesions. Condition 1 introduces transcriptional arrest. Condition 2 combines transcriptional arrest with strengthened boundaries. Condition 3 combines transcriptional arrest, 3.25-fold-strengthened boundaries, and DNA lesions. **c,** Comparison of selected simulation outputs between the baseline state and condition 1 (transcriptional arrest). Relative cohesin occupancy within gene bodies and at CTCF-demarcated boundaries, mean loop length, and boundary-crossing stripe intensity (reflecting one-sided cohesin stalling at CTCF boundaries during loop extrusion) are shown. Points denote independent simulation replicates (n = 20 per condition); horizontal lines denote means. Statistical significance was assessed using a two-sided permutation test on the difference in means (200,000 random label permutations); ***P < 0.001. **d,** Boundary-strength sweep performed in the transcriptionally arrested state to identify a boundary-strength regime that recapitulates UV-associated chromatin features. Fold changes relative to baseline are shown across progressively increasing boundary strengths for cohesin occupancy at CTCF-demarcated boundaries, TAD corner dots, boundary-passing events, and boundary-crossing stripe intensity. **e,** Schematic of lesion classes used in the simulations and their cohesin-blocking capacities during the pre-repair and repair stages. **f,** Two-dimensional parameter sweep varying the duration of the pre-repair and repair phases in simulations with transcriptional arrest and 3.25-fold-strengthened boundaries. Heatmaps show mean loop length as a function of lesion density and phase duration. **g,** Zoomed view of the genome-wide distance-decay profiles shown in Fig. 1b, restricted to genomic separations of up to 2 Mb. Plots show the fold change in contact probability relative to pre-UV WT cells as a function of genomic distance. **h,** Conceptual summary of simulation results.

We first simulated the genome-wide UV-induced phenomenon of transcriptional arrest by removing all RNAPII from the model (Fig. 6c). Although the simulations assume persistent transcriptional shutdown throughout the UV response (whereas transcription gradually recovers *in vivo*), this simplification is unlikely to substantially affect the conclusions relevant to the 6 h time frame captured by our Hi-C data, as transcriptional recovery is not complete until approximately 24 h after a 10 J m^-2^ UVC exposure^69^. Consistent with experimental observations^50,51,53,55^, RNAPII depletion increased cohesin occupancy at CTCF-demarcated TAD boundaries while reducing cohesin occupancy within gene bodies. The model further predicted an increase in loop length and stripe intensity, recapitulating key features of the contact maps generated from our re-analysis of RNAPII degron data (Fig. 6a; Extended Data Fig. 7b). These predictions support the RNAPII barrier-removal model introduced above, wherein RNAPII loss permits longer-range cohesin translocation resulting in increased engagement with CTCF-defined boundaries.

Next, we simulated progressive TAD boundary strengthening, a recurrent feature of both UV and DSB responses, to select a biologically plausible value for the lesion-containing model. We increased boundary strength from the default value of ∼12% (the probability that CTCF captures extruding cohesin;^61^) to as much as six-fold higher (Fig. 6d). As expected, stronger boundaries progressively increased cohesin retention at CTCF-demarcated boundaries while reducing boundary bypass events. Correspondingly, TAD corner-dot intensity increased, reflecting more efficient trapping of cohesin at convergent boundaries. Boundary-crossing stripe intensity decreased with increasing boundary strength, contrasting the increased stripe intensity resulting from RNAPII depletion alone in the first condition but consistent with the absence of stripe enrichment after UV irradiation in Q4 (by post-UV TAD strength) WT TADs (the quartile most susceptible to RNAPII loss upon UV irradiation) (Extended Data Fig. 3d). Genome-wide convergently oriented CTCF sites, which frequently demarcate loop anchors and TAD corners, likewise had no stripe enrichment in WT cells following UV irradiation (Extended Data Fig. 10a,b). This pattern is consistent with increased post-UV CTCF occupancy promoting more efficient bilateral cohesin stalling, shifting TADs from a partially to a fully extruded state. The transition point at which stripe intensity began to stabilise, thereby recapitulating the UV-induced phenotype, occurred at 3.25-fold stronger boundaries.

Finally, we incorporated DNA lesions into the simulations. To capture both TC- and GG-NER events, we distinguished between two lesion classes (Fig. 6e). We modelled Type A lesions, which arise within gene bodies, as lesion-stalled RNAPII. Type B lesions constituted all other–that is, non-RNAPII-stalling– lesions. We further distinguished between a pre-repair phase and a repair phase, with the latter encompassing the assembly and residence time of the core NER factors at lesions^70^. Before repair, only Type A lesions could impede cohesin translocation (through stalled RNAPII). During repair, both lesion classes were permitted to impede cohesin (through the presence of repair intermediates). Because the capacity of NER complexes to act as barriers to cohesin has not been experimentally established, we modelled lesion-associated cohesin stalling probabilistically (Extended Data Fig. 9a; Supplementary Methods). Several additional parameters governing lesion dynamics remain poorly constrained experimentally, including the relative frequency of Type A and Type B lesions and the duration of the pre-repair and repair phases. We estimated lesion density from published measurements of UV-induced photoproduct formation^69^ and constrained the relative frequency of Type A lesions by the lower propensity of transcribed strands to accumulate CPDs (^71^; Extended Data Fig. 9b). Furthermore, we constrained repair to complete far more quickly than pre-repair based on previous measurements of NER kinetics, which reported a slow lesion search by XPC relative to subsequent repair steps^70^. We then performed parameter sweeps across these variables using mean loop length as the optimisation criterion (Fig. 6f; Extended Data Fig. 9a,b). We selected parameter combinations that reproduced the approximately 5–10% changes in loop length observed within 1 h of UV irradiation. This approach identified a high cohesin-blocking probability of 99.3% per 2-second simulation step, a 20% probability that lesions arising within gene bodies were Type A, and pre-repair and repair phases of 40 and 6 min, respectively (Extended Data Fig. 9a,b; Supplementary Methods).

With these optimised parameters, the simulations revealed that RNAPII removal and lesion-associated cohesin stalling exert opposing effects on loop extrusion. Lesion-associated stalling shortened loops by introducing barriers to cohesin translocation, whereas transcription deactivation, entailing RNAPII degradation, lengthened the loops. Loop length therefore reflects the balance between these two processes over time (Fig. 6g,h). Early after UV irradiation, when lesion burden and RNAPII stalling are high, the loop-shortening effect predominates. As repair progresses, RNAPII degrades and lesions are cleared, shifting the balance towards longer loops. This trajectory closely recapitulated the experimental distance-decay curves, in which short-range contacts are enriched shortly after UV exposure, regress towards the baseline by 3 h, and become depleted at later timepoints (6 h). We have previously reported a similar transition from shorter to longer loops in UV-irradiated HeLa cells, although occurring on an accelerated timescale (potentially due to impaired p53 signalling shortening the window available for repair before cell-cycle progression resumes), with short-range contacts greatly enriched at 12 minutes post-UV compared to pre-UV, but consistently decreasing by 30–60 minutes^4^.

Our repair-deficient cell lines provide an additional context with which to interpret the tradeoff between RNAPII depletion and lesion-associated barriers depicted in Fig. 6h. In *XPC*^-/-^ cells, longer loops became more frequent by 3 h post-UV (Extended Data Fig. 1c). Because TC-NER is expected to resolve most transcription-blocking lesions within the first 2 h^72^, lesion-associated RNAPII stalling should be diminished by 3 h, leading to longer loops. By contrast, loop lengths in WT cells remained closer to baseline at 3 h. In WT cells, ongoing GG-NER may generate repair intermediates that continue to impede cohesin translocation after lesion-stalled RNAPII has been cleared, partially counteracting the loop-lengthening effect of transcriptional shutdown. Consistent with this interpretation, *XPA*^-/-^ cells, which are unable to progress beyond damage recognition, showed persistent depletion of short-range contacts compared to WT cells (Extended Data Fig. 1c). These patterns support a model in which not only lesion-stalled RNAPII but also repair-associated barriers stall cohesin. The molecular identity of such barriers remains unresolved, with direct evidence that NER factors physically impede cohesin translocation still lacking.

*XPC*^-/-^ cells have a genome architecture not readily explainable by the principles governing repair-proficient cells. They have shorter chromatin loops, fewer boundary-crossing events, and stronger stripes (Extended Data Fig. 10a-c) and corner-dot signals than WT cells. The shortened loops are consistent with our PHACE-derived hypothesis that XPC attenuates cohesin unloading, as reduced free-cohesin residence time shortened extrusion lengths *in silico* (Extended Data Fig. 10d). However, the remaining *XPC*^-/-^ phenotypes depend on cohesin reaching and being retained at CTCF boundaries, which is difficult to reconcile with their shorter loops and lower CTCF occupancy. We therefore considered whether cohesin that reaches CTCF boundaries is retained there for longer in *XPC*^-/-^ cells. Cohesin residence time increases substantially upon capture by CTCF^61,73^. Further enhancement of this effect might account for the unique architecture of *XPC*^-/-^ cells. To test this possibility, we varied both the free-cohesin residence time and the residence time of cohesin captured at CTCF-bound sites. A combination of reduced free-cohesin residence (approximately one-fourth that of WT) and prolonged residence of CTCF-captured cohesin (approximately twofold greater than WT) recapitulated all *XPC*^-/-^ phenotypes simultaneously (Extended Data Fig. 10d). The redistribution of XPC from TAD boundaries to DNA lesions following UV irradiation may transiently mimic aspects of XPC deficiency, contributing to the corner-dot strengthening observed in both *XPC*^-/-^ cells and UV-irradiated WT cells. However, unlike XPC deficiency, UV irradiation is accompanied by increased CTCF occupancy at TAD boundaries, favouring fully extruded domains and potentially explaining the greater stripe enrichment in *XPC*^-/-^ cells relative to WT cells.

## Discussion

Previously, we showed that UV irradiation is accompanied by increased chromatin insulation, which is associated with enhanced NER efficiency, and by enrichment of short- to mid-range chromatin contacts^4^. Here, by integrating chromatin profiles from repair-deficient cell lines with loop extrusion simulations, we delineate how XPC distribution and the formation and resolution of lesion-associated barriers mediate these architectural changes (Fig. 7). Before damage, chromatin insulation is restricted by XPC at TAD boundaries and by elongating RNAPII within TADs. *In silico*, XPC inhibits cohesin unloading and limits CTCF-mediated cohesin retention, promoting extensive loop extrusion while keeping the boundaries semi-permeable. XPC also facilitates transcription initiation through its established role as an RNAPII cofactor^57^. In turn, ongoing RNAPII elongation during transcription presents a moving barrier to cohesin, further limiting cohesin accumulation at the boundaries while permitting extrusion–and thereby frequent enhancer-promoter communication–within the TADs. As cohesin becomes trapped by multiple RNAPII molecules throughout the TADs, it participates in short-range boundary-crossing events (Fig. 5c) that keep the TADs in a partly extruded state and potentially contribute to the formation of transcriptional hubs^74–76^.

**Figure 7:**
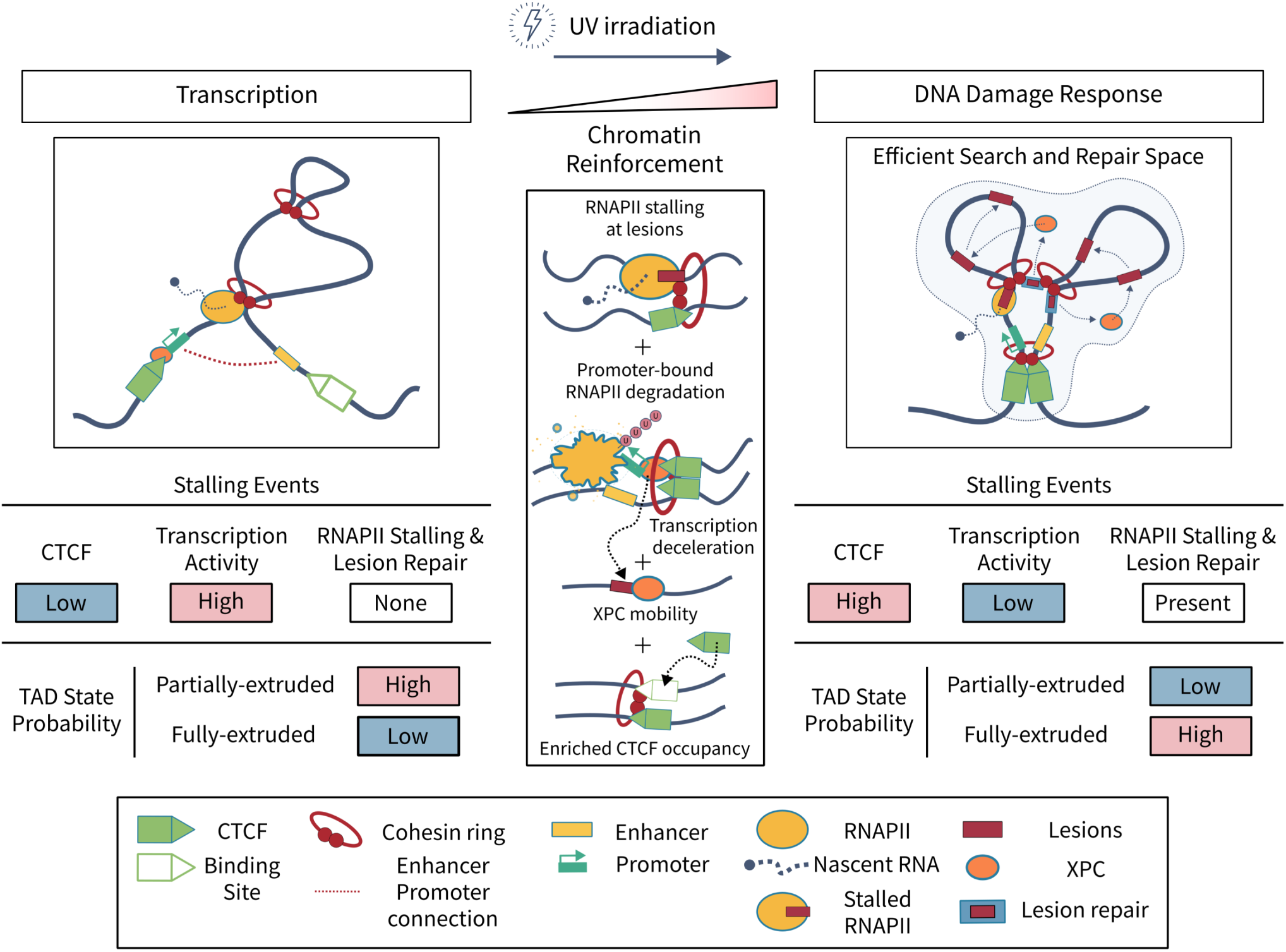
Model of chromatin reorganisation across transcriptional and DNA damage states. Before damage (left panel), XPC occupies TAD boundaries, where it serves as a cofactor of RNAPII, helping assemble the preinitiation machinery and thereby facilitating productive elongation of RNAPII. Elongating RNAPII acts as a moving barrier to cohesin, which is often stalled by CTCF at only one boundary of a TAD, resulting in a partially extruded state. Upon UV-induced damage, several events reconfigure the genome for repair (middle panel). RNAPII becomes stalled at lesions within gene bodies or is degraded; XPC redistributes from TAD boundaries to lesions, and CTCF occupancy increases at TAD boundaries. During repair (right panel), increased CTCF occupancy at TAD boundaries promotes a fully extruded TAD state by constraining cohesin at both boundaries. The reinforcement of TADs partitions damaged chromatin into smaller domains, facilitating lesion recognition by reducing the effective search space. RNAPII stalled at lesions impedes cohesin, shortening loop length. Concurrently, redistribution of the damage sensor XPC from TAD boundaries to lesions promotes repair machinery assembly, constituting an additional barrier to cohesin translocation. The shorter loops bring lesions into closer proximity, reducing the distance over which repair factors must diffuse and thereby facilitating their reuse within the same domain. Counteracting loop shortening, which is most pronounced at early timepoints when lesion burden remains high, RNAPII degradation accompanying lesion resolution and transcriptional shutdown permits cohesin to extrude further.

Upon UV irradiation, the 3D genome reconfigures to prioritise repair over transcription. Loops are shortened by widespread RNAPII stalling at lesions within gene bodies, which converts RNAPII from a moving into a stationary, more effective barrier to cohesin translocation. Repair-associated intermediates constitute an additional class of barriers, capable of constraining cohesin not only within gene bodies but throughout the genome. XPC relocates from TAD boundaries to lesions, transiently mimicking XPC deficiency as predicted *in silico* by increasing cohesin unloading (further shortening loops by limiting the time available for extrusion) and by strengthening CTCF-mediated cohesin retention. The latter combines with the experimentally observed increase in CTCF occupancy at the boundaries to limit inter-TAD contacts. Together with the higher intra-TAD contact frequency (as shorter loops permit more loops per TAD), these changes reinforce TADs after UV, favouring a fully extruded state in which cohesin is constrained at both TAD boundaries. As repair completes and transcription, through global RNAPII degradation, remains shut down, the barriers to loop extrusion are progressively removed, permitting a return towards longer loop lengths.

While we have now extensively characterised the probable mechanisms underlying UV-induced 3D genome reorganisation, its impact on NER efficiency remains largely inferential, supported primarily by correlative evidence and our CTCF and RAD21 depletion experiments. We have now identified the same 3D genome changes (TAD reinforcement and loop shortening), and their association with enhanced repair, in both HeLa and U2OS cells. This convergence supports the idea that these changes serve to bolster NER. Notably, the factors largely driving these changes are themselves NER components: XPC restrains chromatin insulation before damage, and repair-associated intermediates constrain cohesin genome-wide after damage. We therefore interpret these architectural changes not as incidental by-products, but as an integral, actively maintained feature of the repair process. We speculate that the reinforced architecture facilitates both lesion recognition and repair. A pre-partitioned genome may index the damaged chromatin into smaller and more efficient search spaces, as proposed in the context of DSB repair^5,7^ and supported by polymer modelling studies demonstrating accelerated target search within compact chromatin domains^77^. Further, XR-seq studies have demonstrated that repair efficiency is governed not only by lesion identity but also by the chromatin environment in which a lesion resides, resulting in regional domains with shared repair kinetics^4,19^. We further speculate that short loops within highly repaired, active TADs reduce the distance over which NER machinery must diffuse between lesions (Fig. 7). This concept parallels observations from DSB repair, where damaged TADs cluster within shared nuclear territories^8^. The substantially greater lesion burden imposed by UV irradiation (hundreds of thousands at the 20 J m^-2^ dose used here) compared with the sparse DSBs produced in the AsiSI system (∼100 cut sites)^9^, may necessitate organisation at a finer spatial scale to sustain efficient recycling of the repair machinery.

We surmise from our CTCF and RAD21 depletion experiments that when this finer-scale organisation is compromised, NER instead relies on progressively coarser levels of genome organisation. While acute depletion of either factor disrupts TAD organisation and chromatin loops^28,29,78^, only CTCF depletion significantly impaired CPD resolution in our repair synthesis assay. RAD21 depletion alone strengthens compartment segregation, including increasing A-A contacts^15,28–30,78^. Thus, RAD21 depletion reinforces interactions among active chromatin regions within which NER preferentially operates. Consequently, enhanced chromatin compartmentalisation may partially compensate for the loss of loops and TADs by maintaining spatial partitioning of transcriptionally active chromatin, albeit at a substantially larger scale. A similar principle may operate within transcriptional condensates, which cluster active chromatin to locally concentrate transcriptional and regulatory machinery^79^. Although we focused on cis contacts within 10 Mb, future studies might explore how ultra-distal or inter-chromosomal contacts contribute to NER efficiency.

The pre-positioning of XPC at TAD boundaries before damage, consistent with its role as a cofactor of RNAPII at such promoter-enriched regions^80,81^, is compatible with a role in preserving the integrity of these genome architecture-critical elements through both damage prevention and rapid repair. Our PHACE analysis supports a model in which XPC, via RAD23B-mediated turnover of the base excision repair enzyme TDG, promotes completion of DNA demethylation at nearby sites, including the critical 5′ position within the CTCF motif^47,48^. Maintenance of a demethylated state at this position is required for stable CTCF binding, which in turn has been reported to suppress CPD formation^26,82^. After UV irradiation, maintenance of a CTCF-binding permissive state at these sites likely does not rely on canonical TDG-coupled demethylation since XPC redistributes to lesions, TDG is degraded^83^, and transcription–which is coupled to demethylation cycling^84^–shuts down. Alternative mechanisms entailing UV-induced chromatin remodelling, such as the recruitment of CTCF by XPF and XPG^40,41^, may sustain CTCF occupancy at TAD boundaries. Should these regions still incur damage, XPC pre-positioning may accelerate repair by providing local surveillance. Although CTCF binding suppresses CPD formation, it also inhibits NER activity^26,85^. Enrichment of XPC at TAD boundaries could offset this inhibitory effect by enabling immediate damage recognition. Accordingly, we observed enhanced lesion removal at these loci. Beyond local repair, XPC pre-positioning at boundaries may facilitate its rapid redistribution into nearby chromatin domains subject to preferential repair, reinforcing our model that efficient NER proceeds through local rather than long-range search processes.

Our results highlight that nuclear processes do not operate in isolation but act on a shared chromatin substrate, with transcriptional state and repair activity collectively shaping genome organisation. It remains to be determined how additional nuclear processes involved in the DNA damage response, such as DNA methylation cycling, repair machinery assembly and processivity, DNA replication, and cell cycle progression or arrest contribute to the rewiring of chromatin contacts. As genomic technologies continue to advance towards multi-modal and single-cell measurements^86,87^, it will become increasingly feasible to resolve how such processes are coordinated in space and time to maintain genome integrity.

## Methods

### Cell lines

Wild-type U2OS cells, obtained from the American Type Culture Collection and authenticated by short tandem repeat profiling and negative mycoplasma tests, were maintained on 10-cm dishes in low glucose Dulbecco’s Modified Eagle Medium (Gibco, 21885-025) supplemented with 10% (v/v) fetal calf serum (FBS, Gibco, 10500-064) at 37 °C in 5% CO_2_. *XPC*^-/-^ and *XPA*^-/-^ U2OS cell lines were generated by CRISPR/Cas9-mediated deletion as previously described^25^.

### UV irradiation for genomics assays

Cells were irradiated with UVC light (254 nm) using a custom-built UV irradiation setup. Prior to irradiation, culture medium was removed and cells were washed once with PBS and irradiated without medium at room temperature. UV dose (20 J m^-2^) was determined using a calibrated UV radiometer placed at the level of the cell monolayer. Immediately after irradiation, fresh culture medium was added and cells were returned to 37 °C for the indicated recovery times.

### Hi-C library preparation and sequencing

Hi-C libraries were prepared using the Arima Hi-C+ kit (A51008-ARI) and Arima Library Prep Kit V2 (A303011-ARI) according to the manufacturer’s instructions, with the following modifications. Three 10-cm dishes of cells were cultured per library, and cells were quantified after trypsinization to obtain 10 million cells per library. Cells were harvested either before UV treatment or after exposure to 20 J m^-2^ UVC and collected 1, 3, or 6 h post-irradiation.

Chromatin was fragmented by sonication using a BioRuptor Plus (Diagenode) in 1.5-ml sonication tubes for 12 cycles (30 s on/30 s off). AMPure XP beads (Beckman Coulter, A63882) were used for all DNA size-selection steps. For the final purification, libraries were subjected to two sequential AMPure bead cleanups to minimise primer contamination. In the first cleanup, DNA was eluted in 30 µl of water. For re-purification, 29 µl AMPure XP beads were added to the eluate (to account for volume loss during elution), and libraries were eluted in 15 µl of deionized water following washing and drying. All bead incubation steps were performed for 15–20 min to maximise yield.

Final library concentrations and fragment size distributions were assessed using an Agilent TapeStation D1000. Libraries were pooled at equimolar concentrations and sequenced on an Illumina NovaSeq X Plus platform to generate 150-bp paired-end reads. Reads from the initial sequencing run were downsampled to 5 million paired-end reads per library using seqtk (seed = 1) and evaluated with the Arima shallow sequencing QC pipeline. All libraries passed QC and were subsequently re-sequenced on additional NovaSeq lanes to achieve a minimum depth of 550 million read pairs per library. Base calling was performed using Illumina’s standard pipeline (Real-Time Analysis followed by bcl2fastq).

### Hi-C data preprocessing and quality control

Hi-C sequencing reads were processed using Juicer (v2, CPU implementation;^88^) and aligned to the human reference genome (GRCh38). Both intra- and inter-chromosomal contacts were retained for initial quality-control analyses, including assessment of cis/trans contact proportions. Only chromosomes 1 to 22 and X were processed and carried forward to downstream analysis. Juicer quality-control metrics are available in Supplementary Table, S1.

.hic files were generated with Juicer and converted to multi-resolution cooler (.mcool) formats using the hicConvertFormat utility from HiCExplorer^89^ to enable downstream analyses with cooltools (v.0.7.1,^90^). Technical replicates were merged prior to downstream analysis. Contact matrices were iteratively corrected using ICE balancing implemented in cooler (v0.10.4,^91^), which balances contact matrices to remove systematic biases^92^. Copy-number-aware ICE normalisation was additionally applied using NeoLoopFinder^93^, resulting in .mcool files containing both standard ICE-normalised (‘weight’) and CNV-corrected ICE weight columns (‘sweight’, used for all downstream analyses). The effective resolution of each Hi-C library was estimated using the Juicer calculate_map_resolution.sh script, which defines the highest usable resolution as that at which at least 80% of genomic bins contain more than 1,000 contacts. Based on this criterion, library resolutions ranged from 5 to 10 kb. All subsequent analyses used 10 kb resolution, unless stated otherwise.

Replicate concordance was assessed using two complementary approaches. Stratum-adjusted correlation coefficients (SCCs) were computed using HiCRep v0.2.6^94^, which measures similarity between Hi-C contact maps while accounting for the genomic distance dependence of contact frequencies. In parallel, distance-dependent decay profiles were computed from cis contact matrices using the expected_cis method in cooltools, stratifying contacts by genomic distance.

Trans contact enrichment for each sample was assessed using chromosome_matrix method in GENOVA (v1.0.1;^95^) by comparing observed chromosome-by-chromosome contacts from balanced contact maps at 1 Mb resolution to expected contact frequencies estimated from chromosome-level contact signal. Global cis/trans proportions were calculated from the balanced matrices as the fraction of total contact signal assigned to intra- or inter-chromosomal contacts.

### Chromatin compartment analysis

A/B chromatin compartments were identified by eigenvector decomposition of intra-chromosomal Hi-C contact matrices using the eigs_cis method in cooltools at 100 kb resolution, as described previously^4^. Eigenvectors were phased by correlation with GC content. Compartment strength was quantified using saddle plots computed from observed/expected cis contact matrices, and summarised using the saddle strength metric, which compares interaction enrichment within the same compartment (AA + BB) relative to contacts between compartments (AB + BA). Chromosomes 6, 21, and 22 were excluded from compartment analyses due to uncertainty in compartment assignment, as visual inspection of compartment tracks did not allow reliable discrimination between A and B compartments.

### TAD insulation and domain analysis

TADs and boundary insulation profiles were computed using the insulation method in cooltools on ICE-normalised cis contact matrices at 10 kb resolution, using a sliding window of 500 kb, as described previously^4^. Insulation minima were used to identify boundary positions, and boundary strength was quantified based on the prominence of insulation score minima. Boundaries were considered preserved between conditions if they occurred within ±10 kb. Consecutive boundaries were merged to define TADs, excluding excessively large or small domains (>2.0 Mb and <250 kb).

TAD strength was quantified using domain scores^24^, defined as the ratio of intra-domain contact intensity to inter-domain contact intensity calculated from rescaled Hi-C contact matrix snippets spanning each TAD and flanking regions. TADs were grouped into quartiles based on post-UV TAD strength. For metaprofile analyses of chromatin profiling across TAD quartiles and their constituent loops, loops were assigned to the TADs in which they resided, downsampling to the loop count of the quartile containing the fewest loops.

For analyses in lung-derived fibroblasts, contact maps from IMR90 cells were retrieved from the 4DN Data Portal^66^ (with identifier 4DNFIJTOIGOI, in .mcool format), and TADs were identified as described for U2OS cells.

For analyses in HeLa cells, contact maps from pre-UV and 12 min post-UV conditions (retrieved from^4^) were accessed in .mcool format, and TADs and boundaries were identified as described for U2OS cells.

### Chromatin loop analysis

Chromatin loops were identified from ICE-normalised cis contact matrices using the dots method in cooltools at 10 kb resolution, applying default kernel configurations and significance thresholds as described previously^4,22^. Loops were retained based on false discovery rate (FDR)-corrected significance. Loop anchors were defined as the genomic coordinates of interacting pixels passing significance thresholds. Loops and loop anchors were classified as preserved or condition-specific based on overlap within ±20 kb between conditions. Aggregate loop interaction strength was assessed using pile-up analyses of off-diagonal contact matrix snippets centred on loop pixels. Briefly, loop strength was calculated from loop pileups generated with a 100-kb flank as the ratio of the mean signal in the central 3 × 3 pixels to the mean signal in three background corner windows (7 × 7 pixels at upper left, upper right, and lower right).

### CTCF ChIP-seq library preparation and sequencing

ChIP-seq libraries were generated in biological duplicates, with a pooled input control from a UV-irradiated sample. Up to 3 × 10^6^ cells per replicate were seeded from liquid nitrogen stocks onto 10- cm dishes, grown for 1–2 days to 70–80% confluence, and harvested either before or 3 h after UVC irradiation (20 J m^-2^). Cells (10–14 × 10^6^ per sample) were cross-linked in 1% formaldehyde (Sigma-Aldrich, 252549) for 15 min with agitation, quenched in 0.1 M glycine for 5 min, and collected by centrifugation (800 × g, 10 min, 4 °C). Cell pellets were washed twice in PBS containing 0.5% Igepal (Sigma) and protease inhibitor cocktail (Roche, 11836170001), re-pelleted under the same conditions, snap-frozen in liquid nitrogen and stored at -80 °C until shipment on dry ice to Active Motif.

ChIP-seq processing was performed by Active Motif (Carlsbad, CA). Chromatin was prepared by cell lysis followed by washing and sonication using the PIXUL Multi-Sample Sonicator (Active Motif; catalog no. 53130) to obtain an average DNA fragment size of 200–1,000 bp. Chromatin yield was determined by reverse crosslinking an aliquot at 65 °C, followed by RNase and proteinase K treatment and DNA purification using SPRI beads (Beckman Coulter). DNA concentrations were measured using a Qubit fluorometer (Thermo Fisher Scientific), and total chromatin yield was extrapolated based on the initial chromatin volume. Chromatin aliquots were precleared for immunoprecipitation with protein G agarose beads (Invitrogen) and incubated with an AbFlex anti-CTCF antibody (Active Motif; cat. no. 91285; RRID: AB_3216313; lot no. 25006102-11). After washing, immune complexes were eluted using SDS buffer, treated with RNase and proteinase K, and de-crosslinked by overnight incubation at 65 °C. ChIP DNA was purified by phenol-chloroform extraction followed by ethanol precipitation. To repair UV-induced DNA lesions that can inhibit PCR amplification, UV-irradiated ChIP and input samples were treated with the PreCR Repair Mix (New England BioLabs, M0309S) according to the manufacturer’s protocol. Sequencing libraries were prepared using the NEB DNA Library Prep Kit on an Apollo 342 system (Wafergen Biosystems / Takara) and sequenced on an Illumina NextSeq 2000 platform as 50 bp paired-end reads.

### CTCF ChIP-seq analysis

Base calling and demultiplexing were performed with bcl2fastq v2.20, without adapter trimming during demultiplexing. FASTQ files were processed using the nf-core/chipseq pipeline (v.2.1.0,^96^), with default parameters unless stated otherwise. Adapter trimming, alignment to the human genome (GRCh38/hg38) using Chromap^97^, and peak calling with MACS3^98^ were performed within the pipeline. Signal tracks (bigWig files), normalised such that the total signal summed to one million mapped reads, and peak sets generated by nf-core/ChIP-seq were uploaded to the GEO. For all secondary analyses, signal tracks normalised to reads-per-genomic-content (RPGC) were used (deeptools, v3.5.6,^99^).

To independently assess data quality, ChIP-seq libraries were additionally processed using the ENCODE ChIP-seq pipeline following established quality control and reproducibility guidelines^100^. FRiP scores ranged from 0.18 to 0.46 across libraries, and cross-correlation metrics met ENCODE quality thresholds (NSC ≥ 1.1, RSC ≥ 0.8). Replicate concordance was confirmed by IDR analysis^101^, with rescue ratios of 1.05–1.93 and self-consistency ratios of 1.03–1.23, all within ENCODE-recommended limits.

Peak calling with MACS3 yielded approximately 61,000–76,000 peaks per replicate. Of these, 49,800– 60,000 peaks per replicate met ≥5-fold enrichment and FDR ≤ 5%. Reproducible CTCF peaks were defined as sites whose summits in biological replicates were within 150 bp of each other. Fixed 300 bp windows centred on the mean summit position were used for downstream analyses, yielding 47,000– 62,000 reproducible peaks per condition.

### ATAC-seq and damage-seq data processing and analysis

ATAC-seq and damage-seq, as well as ChIP-seq (XPC, H3K4me1/3, H3K27ac) used in this study were generated previously^25,43^. For the integrative analyses performed here, datasets were reprocessed or re-normalised as needed to ensure consistent normalisation and comparability across assays, replicates, and conditions. ATAC-seq data were reprocessed using the nf-core/atacseq pipeline (v2.1.2;^96^), and signal tracks were normalised using reads-per-genomic-content (RPGC) (deeptools, v3.5.6^99^). Standard filtering steps were applied, including removal of low mapping quality reads.

For data deposition, legacy fold-change-normalised ATAC-seq signal tracks from *XPC*^-/-^ cells, generated previously but not included in earlier work, were released to maintain direct comparability with previously published WT fold-change tracks^25^.

Damage-seq data were reprocessed using the XR-seq/Damage-seq pipeline (v1.0), incorporating adapter trimming, alignment, filtering, and simulation, available at CompGenomeLab/xr-ds-seq-snakemake^68^. In brief, reads were used to define exact damage-site coordinates, which were then mapped to 1 kb intervals across the human genome and normalised to reads per million (RPM). We also generated synthetic NGS reads using the simulation tool Boquila^102^ to assess expected damage profiles based on the (di-)nucleotide frequency distribution observed in the experimental data. Simulation data (expected damage-seq) were also RPM normalised.

### Integrative analyses

Integrative analyses were performed by intersecting Hi-C-derived features (TADs, loop anchors, and compartments) with ChIP-seq, ATAC-seq, and damage-seq signal tracks. Signals were aggregated over defined genomic features and compared across genotypes and conditions using custom scripts. For inferential comparisons in the integrative genomics analyses, two-sided nonparametric tests were used as specified in the corresponding figure legends. No formal normality tests were performed. Multiple-testing control applied during genomic feature calling, including FDR-based loop and ChIP-seq peak calling, is described separately. Test statistics are provided in Supplementary Table S11.

For permutation analyses, we tested whether XPC sites were positioned closer to CTCF than expected under a chromatin accessibility-matched null model. XPC summits were scored for local ATAC-seq intensity and grouped into accessibility quantile bins, and the same ATAC-seq signal thresholds were used to partition an ATAC-defined genomic universe restricted to chr1–22 and chrX. In each permutation, XPC summits were shu ed within accessible genomic regions assigned to the same ATAC-intensity bin, thereby preserving both chromosome identity and the overall accessibility distribution of the observed XPC set. For the observed and permuted datasets, distances from each XPC summit to the nearest CTCF summit were then computed.

To evaluate the association between XPC occupancy and chromatin insulation within wild-type cells, genome-wide covariate-matched analyses were performed in genomic bins with focal (high XPC) or matched control (low XPC signal) matched for RNAPII, ATAC-seq or histone mark (H3K4me1/3, H3K27ac) signal, and insulation scores and CTCF ChIP-seq signal were compared between focal and matched bin sets. Match quality was assessed using standardized mean differences, and analyses were repeated after progressively restricting the focal set to the best-matched high-XPC bins to minimise residual covariate imbalance. Covariate matching was performed using the nullranges R package (v1.17.3,^103^), using the matchRanges method.

For analyses within lung-derived fibroblasts, RNAPII pre- and post-UV ChIP-seq data from MRC5 cells were retrieved (from^58^) and projected onto TAD strength-grouped quartiles from IMR90 cells^66^.

For analyses within HeLa cells, RNA-seq data (retrieved from^4^) were processed using the nf-core/rnaseq pipeline (v3.23.0;^96^) with default parameters, and alignments were CPM-normalised to generate signal tracks. The RNAPII ChIP-seq signal track for HeLa cells was retrieved from the ENCODE data portal^104^ under the identifier ENCFF144IVU. First-12-min CPD repair XR-seq data for HeLa cells (from^68^, in FASTQ format, NCBI-SRA PRJNA608124) were processed using the same XR-seq/Damage-seq pipeline (as for U2OS damage-seq), as previously described^4^. Briefly, reads were used to define exact repair-site coordinates, which were then mapped to 1 kb intervals across the human genome and normalised to reads per million (RPM).

### Phylogenetic analysis of co-evolution with PHACE

The Phylogeny-Aware Detection of Molecular Coevolution (PHACE;^44^) workflow was used to assess coordinated amino-acid substitutions between proteins. PHACE detects correlated changes between pairs of amino-acid positions by tracing evolutionarily independent substitution events. Because accurate inference of coevolution depends on correct reconstruction of evolutionary relationships, PHACE was applied to concatenated multiple sequence alignments (MSAs) of orthologous sequences from the proteins under comparison, with paralogous sequences excluded. To this end, an orthology assignment workflow was implemented. A diverse set of 174 eukaryotic species (Supplementary Table, S2) was used for orthology assignment of 45 proteins involved in GG-NER and in chromatin architecture (UniProt IDs in Supplementary Table, S3). Homologous sequences were identified by three iterations of PSI-BLAST (BLAST+ v2.16.0)^105^ using human proteins as queries. Sequences were aligned with MAFFT (v7.526)^106^ using the E-INS-i algorithm and trimmed with ClipKIT (v2.4.1)^107^ using the kpic-smart-gap option. For each trimmed MSA, maximum-likelihood (ML) trees were reconstructed with IQ-TREE (v3.0.1)^108^ using the best-fitting substitution model selected by ModelFinder^109^ from a subset of general models (WAG, JTT, LG, and Q.Pfam), with 1,000 ultrafast bootstrap replicates for branch support^110^. To strengthen orthology assignment, lineage information was integrated to the tree through the ETE3 module (v3.1.2)^111^, and domain architecture of sequences was assessed using hmmscan from HMMER (v3.4)^112^ against the Pfam-A database (release 38.0)^113^. Orthologs of each human query sequence were selected through manual evaluation of species distribution, duplication patterns, shared domain architecture and consistency with the literature (See Supplementary Table, S4 for selected orthologs). For each query protein, a single orthologue per species was retained by selecting the sequence phylogenetically closest to the human query. The resulting orthologous sequences were realigned with MAFFT E-INS-i and trimmed to positions present in the human sequence. Best-fitting evolutionary models were then re-estimated from the same model subset using ModelFinder, using only the assigned orthologs for each protein. Trimmed MSAs were concatenated into multi-protein alignments, and species lacking an ortholog for a given protein were represented by gap-only blocks for that partition to preserve alignment length and partition structure across species. Because this required explicit handling of artificial gap blocks, the PHACE workflow was adapted so that branch-level contributions from gap-only blocks were excluded from the coevolution score, reducing artefactual signal from missing-protein partitions. A partitioned ML tree search^114^, followed by ancestral sequence reconstruction (ASR), was then performed in IQ-TREE using protein-specific models, and PHACE was run on the resulting multi-protein MSA, ML tree, and ASR outputs. For the pairwise XPC-CTCF analysis, the orthology assignment workflow was first applied to a curated set of 508 metazoan species (Supplementary Table, S5). Species without an assigned ortholog for either XPC or CTCF were then excluded, leaving 431 species (See Supplementary Table, S6 for selected orthologs). The same PHACE input-generation workflow, including concatenation of trimmed MSAs, partitioned ML tree search, and ASR using protein-specific models, was then applied to the ortholog sequences from this retained species set. Accordingly, no artificial missing-protein gap blocks were present in the two-protein MSA; in the absence of such blocks, the modified PHACE workflow is equivalent to the original PHACE implementation.

### siRNA transfection and knockdown validation

For siRNA-mediated protein depletion, U2OS cells were seeded in six-well plates at 4 × 10⁵ cells per well and 3 mL DMEM supplemented with GlutaMAX and 10% FBS. The following day, medium was replaced with fresh medium and cells were transfected using Lipofectamine RNAiMAX. For each transfection, siRNA was diluted in 500 µl Opti-MEM and mixed with 5 µl RNAiMAX, with siRNAs used at a final concentration of 20 nM. Twenty-four hours after transfection, cells were trypsinised and reseeded onto 13-mm glass coverslips in six-well plates containing 2 mL medium per well. Cells were grown to approximately 80% confluency before UVC irradiation (254 nm, 100 J m-2) and downstream immunofluorescence analyses. The following siRNAs were used: non-targeting control siRNA, siRAD21, siCTCF1, siCTCF2, siDDB2 and siXPC. siRNA sequences and suppliers are listed in Supplementary Table S7. Knockdown efficiency was assessed in parallel siRNA-transfected cells harvested 48 h after transfection by immunoblotting and RT-qPCR. For immunoblotting, cells were lysed in NETN lysis buffer containing 100 mM NaCl, 20 mM Tris-HCl pH 8.0, 0.5 mM EDTA and 0.5% NP-40, freshly supplemented with EDTA-free protease inhibitor cocktail, 0.1% benzonase and 1 mM MgCl₂. Lysates were incubated on ice for 30 min, sonicated for three cycles of 30 s on/30 s off using a Bioruptor Plus sonication device and clarified by centrifugation at 16,000g for 15 min at 4°C. Protein concentrations were determined using Bradford reagent. Equal amounts of total protein were mixed with 4× Laemmli sample buffer containing 10% β-mercaptoethanol, denatured at 95°C for 5 min, separated by 10% SDS-PAGE and transferred to nitrocellulose membranes using a Trans-Blot SD semi-dry transfer system. Membranes were blocked in 5% non-fat dry milk in PBS-T for 1 h at room temperature, incubated with primary antibodies overnight at 4°C, washed three times in PBS-T, incubated with peroxidase-conjugated secondary antibodies for 1 h at room temperature and washed again three times in PBS-T. Protein bands were detected using enhanced chemiluminescence and imaged using a ChemiDoc MP system. Antibodies used for immunoblotting, including suppliers and dilutions, are listed in Supplementary Table S8. For RT-qPCR verification of knockdown efficiency, total RNA was extracted using the RNeasy Mini Kit (Qiagen) according to the manufacturer’s instructions. RNA concentration and quality were assessed using a NanoDrop 2000c spectrophotometer. cDNA was generated from 1 µg RNA using the iScript cDNA Synthesis Kit (Bio-Rad). Quantitative PCR reactions were performed in duplicate using the KAPA SYBR FAST system (Roche) on a CFX384 Real-Time PCR system with a C1000 Touch Thermal Cycler. Relative RNA levels were calculated using the 2−ΔΔCT method and normalised to GAPDH or B2M. RT-qPCR primers are listed in Supplementary Table S9. Test statistics are provided in Supplementary Table S10.

### DNA repair synthesis assay

DNA repair synthesis was measured as described previously^25^. Briefly, control or siRNA-transfected U2OS cells were grown on 13-mm glass coverslips to ∼80% confluency and subjected to localised UVC irradiation (254 nm, 100 J m^-2^) through 5-µm polycarbonate filter membranes to generate discrete nuclear UV damage spots. Following recovery for the indicated times, cells were incubated with 10 µM EdU for 1 h to label sites of repair synthesis. Cells were then pre-extracted, fixed with paraformaldehyde, and DNA was denatured prior to detection of incorporated EdU using Click-iT chemistry according to the manufacturer’s instructions. UV damage was visualised by immunostaining for cyclobutane pyrimidine dimers (CPDs; antibodies listed in Supplementary Table S8). Nuclei were counterstained with DAPI.

Images were acquired on a DMI600 B inverted fluorescence microscope using a 63× oil-immersion objective under identical acquisition settings across all conditions. Repair synthesis was quantified by measuring EdU signal intensity within CPD-positive nuclear spots after background subtraction. S-phase cells displaying high nuclear EdU fluorescence were excluded from the quantification. Representative images consistent with the population-level data were selected from randomly acquired fields of view. Brightness and contrast adjustments were applied uniformly across all samples. Immunofluorescence images were quantified using FIJI (ImageJ, V.2.16.0). Statistical analyses were performed in GraphPad Prism (version 10.16.1, GraphPad Software). Outliers were identified using the ROUT method (Q = 1%) and excluded prior to analysis. Data were normalised within each experiment such that 0% corresponded to background and 100% to the median value of the control group. Three independent biological experiments were analysed. Statistical significance was assessed using one-way ANOVA followed by a two-sided Dunnett’s multiple-comparison test comparing each experimental condition to the control. Dunnett-adjusted P values were reported for post hoc comparisons. No formal test of normality was performed due to the small number of independent biological experiments per condition (n=3). Test statistics are provided in Supplementary Table S10.

### Immunofluorescence detection of GG-NER factor recruitment

Recruitment of GG-NER factors to UV-induced DNA damage was assessed by immunofluorescence as described previously^25^. Control or siRNA-transfected cells grown on coverslips were subjected to localised UVC irradiation through micropore filters as described above, followed by recovery for the indicated times. After pre-extraction and fixation, DNA was denatured and cells were blocked prior to incubation with antibodies against CPDs together with antibodies against the indicated GG-NER factors (XPC, XPB, XPA). Primary and secondary antibodies, suppliers and dilutions are listed in Supplementary Table S8. Following incubation with fluorescent secondary antibodies and DAPI staining, samples were mounted and imaged. Images were acquired on a DMI600 B inverted fluorescence microscope using a 40× objective under identical acquisition settings across all conditions. Recruitment was quantified by measuring protein signal intensity at CPD-positive spots relative to nuclear background in at least 100 cells per condition per independent biological experiment. Representative images consistent with the quantified population-level data were selected from randomly acquired fields of view. Immunofluorescence images were quantified and analysed as described above, including ROUT outlier exclusion (Q = 1%), within-experiment normalisation, and statistical analysis by one-way ANOVA followed by a two-sided Dunnett’s multiple comparison test comparing each experimental condition to the control. Dunnett-adjusted P values were reported for post hoc comparisons. No formal test of normality was performed due to the small number of independent biological experiments per GG-NER factor (n=3). Test statistics are provided in Supplementary Table S10.

### Software

Sequencing-data analyses were conducted using Python version 3.12 and R version 4.5.2 with Bioconductor version 3.22. Microscopy images were quantified using FIJI and ImageJ as described above, with statistical analyses performed in GraphPad Prism. The manuscript was written and edited collaboratively on GitHub using Manubot^115^. Plots were generated with seaborn version 0.13.2^116^, and figures were assembled using the open-source vector graphics software Inkscape.

## Acknowledgements

O.A. discloses support by TUBITAK (The Scientific and Technological Research Council of Türkiye) 2232 (Grant ID: 118C320) International Fellowship for Outstanding Researchers Program, Young Scientist Grant (BAGEP) by Science Academy (Türkiye), and European Molecular Biology Organisation Installation Grant. H.N. is supported by the Swiss National Science Foundation grant 219340. M.N.Y. discloses support for the research of this work from the Novartis Foundation for Medical-Biological Research, Young Investigator Grant (#24C191) and from the Graduate Research Campus (University of Zurich, 2024 Career Grant). Hi-C libraries were sequenced at the Functional Genomics Center Zurich. CTCF ChIP-seq experiments were performed and sequenced by Active Motif. Computational analyses were supported by access to high-performance computing resources provided by Science IT at the University of Zurich and toSUn cluster at Sabancı University. We thank Dr. Tobias Widmer for comments on the manuscript and Dr. Nurdan Kuru for feedback on the co-evolution analysis.

## Author Information

V.O.K. and M.N.Y. wrote the first draft, performed initial analyses, and interpreted the data, including extensive exploration of alternative mechanisms. V.O.K. led and performed the primary computational analyses, including simulations, drove their conceptual development, and prepared and refined the figures. O.A. oversaw application of PHACE tools. M.M. curated eukaryotic datasets and optimised and implemented multi-protein PHACE analysis. L.T. performed non-genomics assays. H.N. oversaw the non-genomics experiments. M.N.Y. initiated the collaboration, generated Hi-C libraries, and coordinated CTCF ChIP-seq experiments. V.O.K., O.A., H.N., and M.N.Y. contributed to experimental design. M.N.Y. and H.N. conceived the project and acquired consumables funding. All authors discussed the results and contributed to manuscript editing.

## Additional Information

Supplementary Information is available for this paper. Correspondence and requests for materials should be addressed to M.N.Y.

## Supplementary information

### Supplementary Tables

Hi-C quality-control metrics, PHACE datasets, and reagents used in functional assays. S1: Quality-control metrics for all Hi-C libraries, computed using Juicer^88^. S2: Eukaryotic species used in the global PHACE analysis. S3: UniProt identifiers for proteins included in the PHACE analysis. S4: Sequence identifiers of manually assigned orthologs of 45 proteins across 174 eukaryotic species. S5: Metazoan species included in the PHACE analysis of XPC-CTCF co-evolution. S6: Sequence identifiers of manually assigned orthologs of XPC and CTCF proteins across 431 metazoan species. S7: Antibodies used for immunofluorescence and immuno (Western) blotting, including target, host species, supplier, application and dilution. S8: siRNA reagents used for protein depletion, including target sequence and supplier. S9: RT-qPCR primers used for knockdown validation, including sequence and target gene. S10: Statistical tests for non-genomics assays. S11: Statistical tests for integrative analyses.

### Supplementary Methods

Detailed description of one-dimensional stochastic modelling of loop extrusion, transcription and nucleotide-excision repair.

## Data Availability

Raw and processed Hi-C, ATAC-seq (*XPC*^-/-^), and CTCF ChIP-seq generated or first released in this study have been deposited in the NCBI Gene Expression Omnibus (GEO,^117^) under accession numbers GSE318309, GSE318235, and GSE318236 and will be released upon publication. Previously published datasets analysed in this study are available under BioProject PRJNA872306 (GSE227009), GSE261073, and GSE260917.

## Code Availability

The code for the analyses presented in this study is available publicly at the following repository: CompGenomeLab/ggner-3d. Simulation engine for transcription, nucleotide-excision repair, and loop extrusion is available publicly at CompGenomeLab/ner_rnapii_3d. PHACE implementation for multi-protein co-evolution analysis is available publicly at CompGenomeLab/PHACE-MP.

## Extended Data Figures

**Extended Data Fig. 1.**
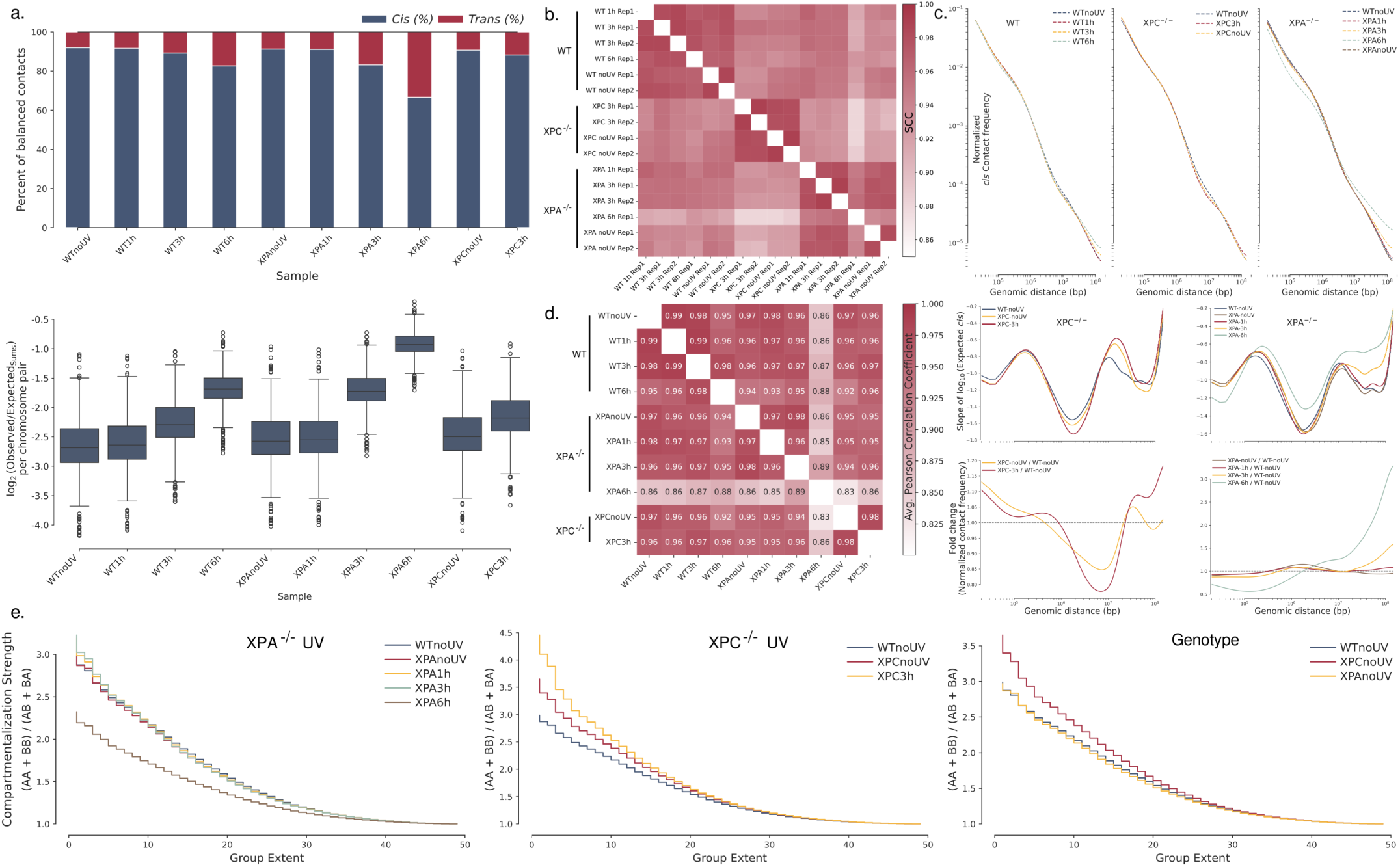
Hi-C data quality, concordance, and compartment organisation across genotypes and timepoints. **a,** Global contact distributions across Hi-C libraries from wild-type (WT), *XPC*^-/-^, and *XPA*^-/-^ U2OS cells before and after UV irradiation. Proportions of cis and trans contacts are depicted as barplots. Distributions of normalised Hi-C contact frequencies calculated for each chromosome-chromosome pair are shown as boxplots including median and interquartile range, with whiskers extending to 1.5x the interquartile range. **b,** Pairwise stratum-adjusted correlation coefficients (SCC) calculated from genome-wide cis Hi-C contacts across all libraries, showing concordance among biological replicates and across genotypes and timepoints. **c,** Distance-decay profiles derived from ICE-normalised Hi-C contact maps at 10 kb resolution for WT, *XPC*^-/-^, and *XPA*^-/-^ U2OS cells across UV irradiation timepoints. Top, genome-wide average cis contact probability as a function of genomic distance. Middle, slope of the distance-decay curves. Bottom, fold change in normalised cis contact frequency relative to the pre-UV condition within each genotype. **d,** Pairwise Pearson correlations (r) of compartment eigenvectors (PC1) computed from ICE-normalised Hi-C maps at 10 kb resolution are shown as a heatmap across all conditions. Eigenvector orientation was standardised per chromosome prior to comparison. **e,** Compartment strength profiles from WT, *XPC*^-/-^, and *XPA*^-/-^ cells across UV irradiation timepoints, showing normalised enrichment of same-compartment over different-compartment contacts (AA + BB)/(AB + BA) for loci up to 100 Mb apart, plotted as a function of compartment rank (group extent), where smaller extent corresponds to bins with stronger compartment identity. Quality-control metrics for all Hi-C libraries are provided in Supplementary Table S1.

**Extended Data Fig. 2.**
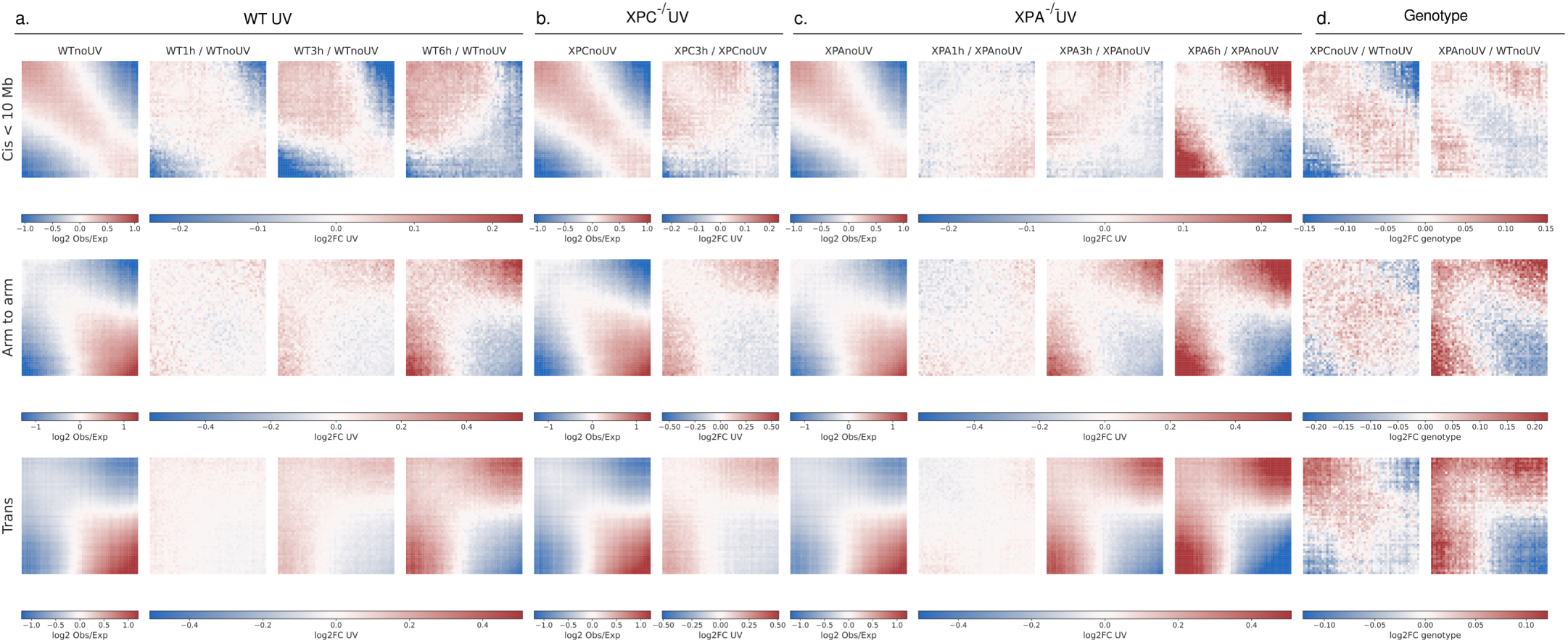
Compartment organisation across genomic scales, genotypes, and UV exposure timepoints. **a-c,** Compartment saddle plots derived from ICE-normalised Hi-C maps for wild-type (WT), *XPC*^-/-^, and *XPA*^-/-^ U2OS cells before and after UV irradiation. For each condition, contacts are shown at three genomic scales: same-chromosomal arm cis (<10 Mb), arm-to-arm (∼100-250 Mb), and trans. Axes are ordered by compartment eigenvector (PC1) value, with quadrants corresponding to B-B, B-A, A-B, and A-A contacts. Within each genotype, the basal (non-UV) condition is shown alongside UV-induced changes relative to baseline. Colour scales are matched within each scale and genotype across timepoints. **d,** Cross-genotype comparisons of compartment contacts in untreated cells, shown as saddle difference maps between *XPC*^-/-^ or *XPA*^-/-^ and WT. Colour scales are matched across genotype comparisons for each interaction class.

**Extended Data Fig. 3.**
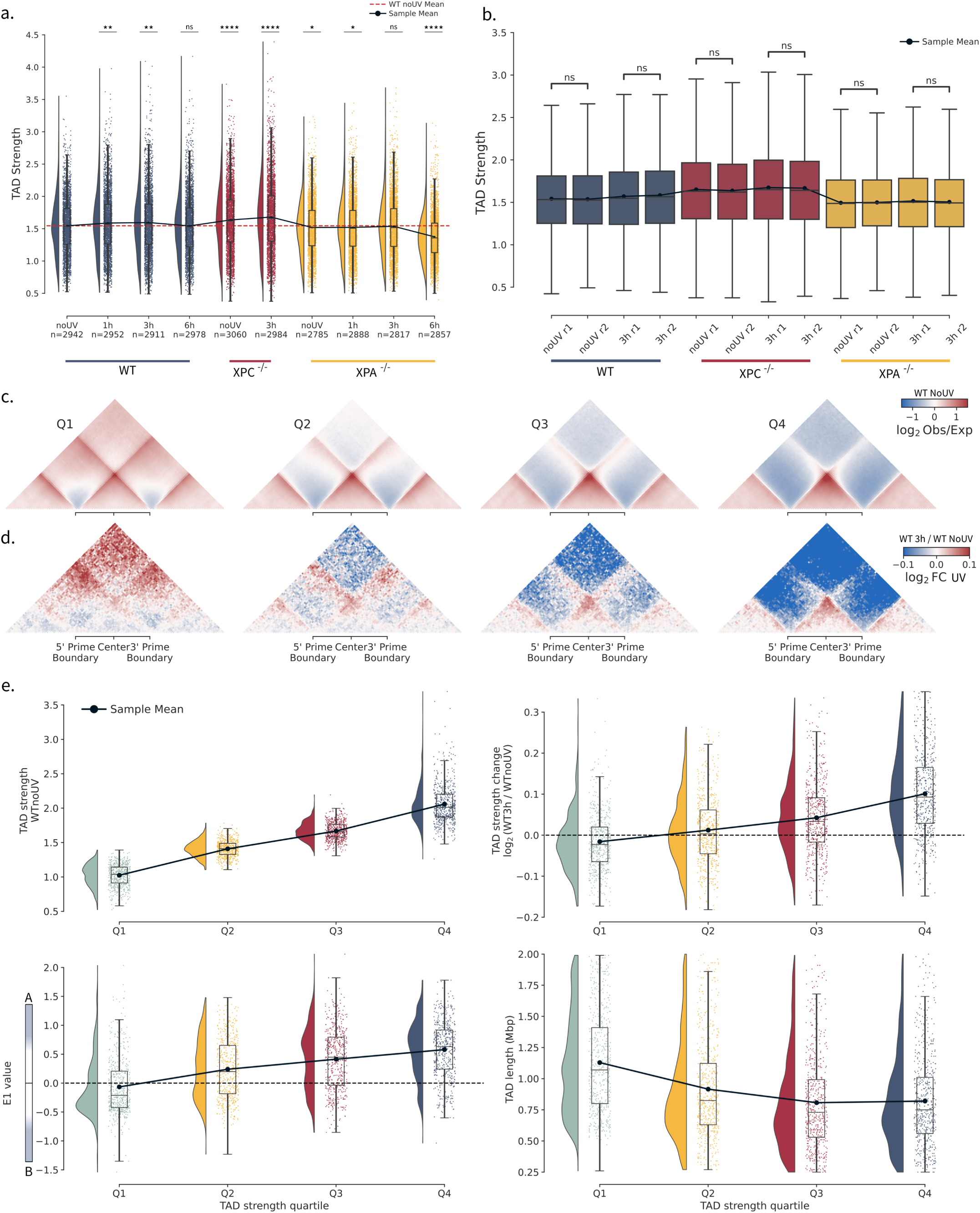
Domain- and loop-level chromatin organisation across genotypes and conditions. **a,** Distributions of topologically associating domain (TAD) strength across wild-type (WT), *XPC*^-/-^, and *XPA*^-/-^ U2OS cells under indicated UV conditions. TAD strength was quantified using domain scores^24^, defined as the ratio of intra-domain contact intensity to inter-domain contact intensity calculated from rescaled Hi-C contact matrix snippets spanning each TAD and flanking regions. Violin plots show per-TAD strength distributions, with boxplots indicating median and interquartile range, whiskers extending to 1.5× the interquartile range, and lines denoting sample means. The dashed horizontal line denotes the mean TAD strength in WT untreated cells. Individual data points denote TADs (WT: noUV, n = 2,942; 1 h, n = 2,952; 3 h, n = 2,911; 6 h, n = 2,978; *XPC*^-/-^: noUV, n = 3,060; 3 h, n = 2,984; *XPA*^-/-^: noUV, n = 2,785; 1 h, n = 2,888; 3 h, n = 2,817; 6 h, n = 2,857). Statistical comparisons were performed against the WT untreated condition with a two-sided Mann-Whitney U test. *P < 0.05; **P < 0.01; ***P < 0.001; ****P < 0.0001; ns, not significant. **b,** TAD strength across independent biological replicates (r1 and r2) for each genotype and condition, evaluated over the TADs of WT noUV cells (n = 2942). Boxplots show median and interquartile range of TAD-strength values within each replicate. Whiskers extend to 1.5× the interquartile range. Points and lines denote replicate means. Statistical significance between replicate TAD-strength distributions was assessed using a two-sided Mann-Whitney U test; ns, not significant. **c,** Aggregate Hi-C contact maps centred on TADs from undamaged WT cells, stratified by 3 h post-UV TAD strength. Maps show observed / expected contact frequencies. **d,** Aggregate Hi-C contact maps showing UV-induced changes in TAD contact structure across quartiles. Difference maps (WT 3 h / WT noUV; log2 fold change) are centred on TADs as in c. **e,** Distributions of TAD strength in WT cells prior to UV irradiation (top left), change in TAD strength 3 h after UV (top right), compartment eigenvector (E1) signal (bottom left), and TAD length (bottom right), shown across TAD quartiles grouped by 3 h post-UV TAD strength. Violin plots show data distributions; boxplots, median and interquartile range; points, individual TADs.

**Extended Data Fig. 4.**
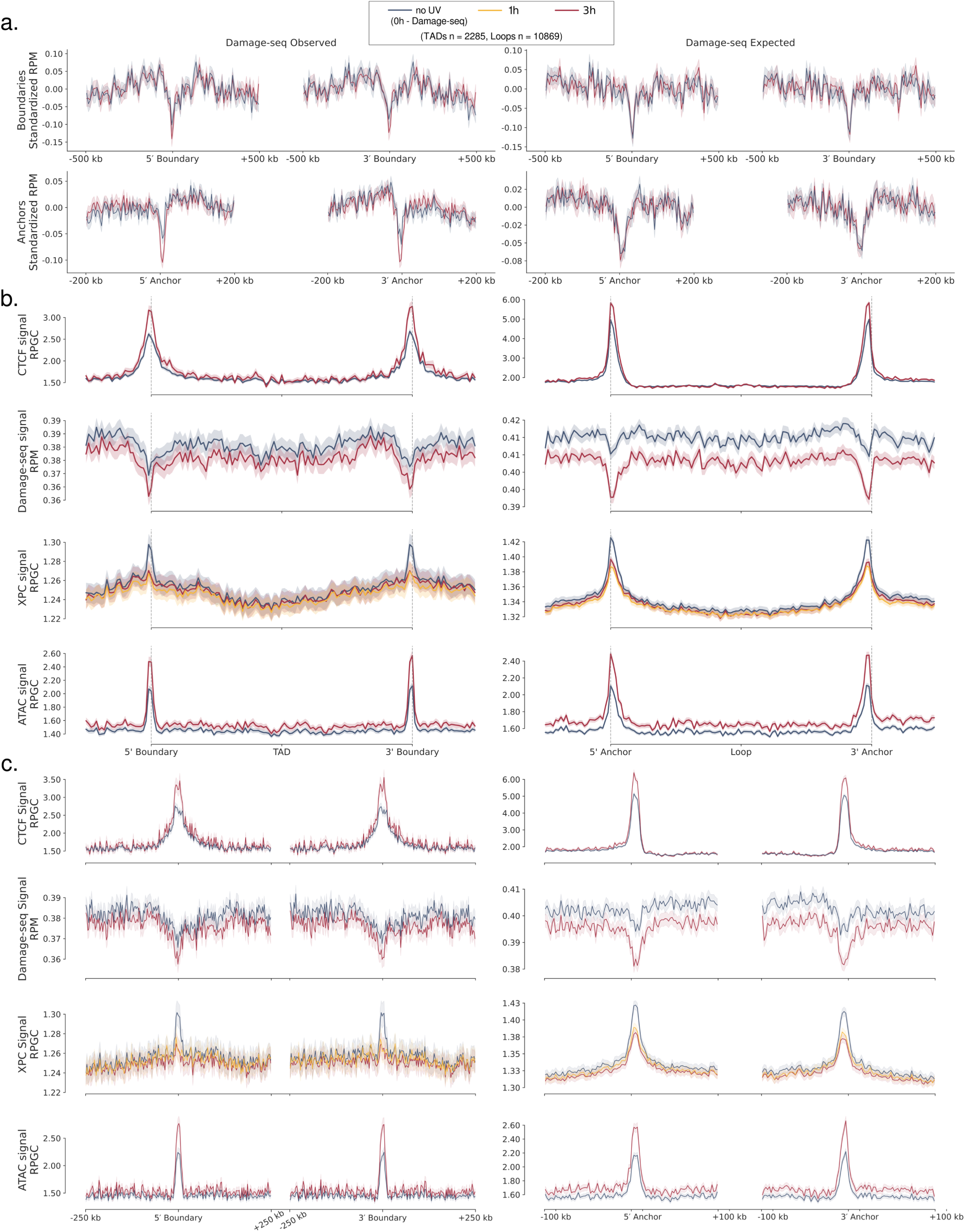
Chromatin features across unstratified TADs and chromatin loops. **a,** Experimental (left column) and simulated (right column) damage-seq signals across topologically associating domain (TAD) boundaries (top row) and loop anchors (bottom row). Cyclobutane pyrimidine dimer (CPD) damage-sequencing (damage-seq) signals were library-size normalised (RPM) and z-normalised to highlight signal changes at feature centres relative to flanking regions. The simulated damage-seq signal reflects sequence-context-derived damage susceptibility and provides an expected baseline for comparison with experimental damage profiles. **b,** Metaprofiles of CTCF occupancy (ChIP-seq), CPD damage (damage-seq), XPC occupancy (ChIP-seq), and chromatin accessibility (ATAC-seq) across TADs (left column) and chromatin loops (right column) preserved in WT cells from before to 3 h after UV. Dashed vertical lines denote TAD boundaries and loop anchors. **c,** The same features, aligned to 5’ and 3’ TAD boundaries and loop anchors. Line plots show mean signal across regions; shaded bands, mean ± SEM.

**Extended Data Fig. 5.**
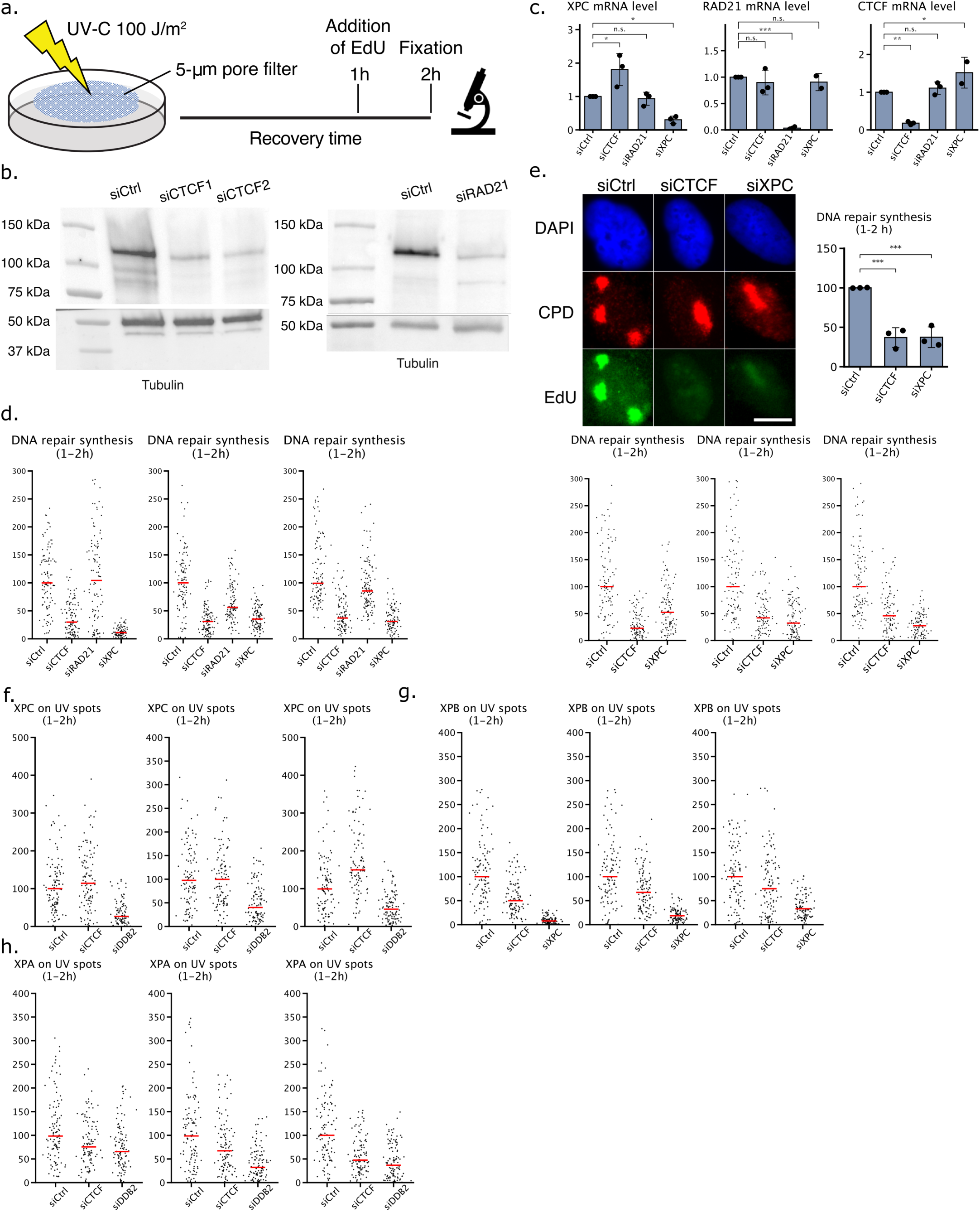
Repair synthesis and repair factor recruitment at local CPD lesions. **a,** Schematic of repair synthesis assay, adapted from^25^. **b,** Immunoblot analysis of knockdown efficiency for RAD21 and two independent CTCF siRNA duplexes, including the second duplex featured in **e,** with tubulin as a loading control. **c,** RT-qPCR measurements of CTCF, RAD21 and XPC mRNA following siRNA treatment, confirming effective target depletion and showing that XPC mRNA is not reduced by siCTCF. Dots denote independent biological experiments (n = 3). Bars denote means and error bars denote SD. Statistical significance was assessed using one-way ANOVA followed by a two-sided Dunnett’s multiple-comparison test comparing each experimental condition to the control. Significance: n.s., P ≥ 0.05; *P < 0.05; **P < 0.01; ***P < 0.001. **d,** DNA repair synthesis assay measured by EdU incorporation at cyclobutane pyrimidine dimer (CPD) foci induced by local UV irradiation through a 5-µm pore filter, quantified at 1–2 h after UV. **e,** DNA repair synthesis as in **d,** using an independent siCTCF duplex. Dots denote independent biological experiments (n = 3). Bars denote means and error bars denote SD. Representative images (DAPI, CPD, EdU) are shown; scale bars, 10 µm. Statistical significance was assessed with one-way ANOVA followed by a two-sided Dunnett’s multiple comparison test comparing each experimental condition to the control. ***P < 0.001. f-h, Immunofluorescence quantification of XPC, XPB and XPA enrichment at CPD foci 1–2 h after UV in the indicated knockdown conditions. Data are shown as per-experiment dot plots, n=3 experiments.

**Extended Data Fig. 6.**
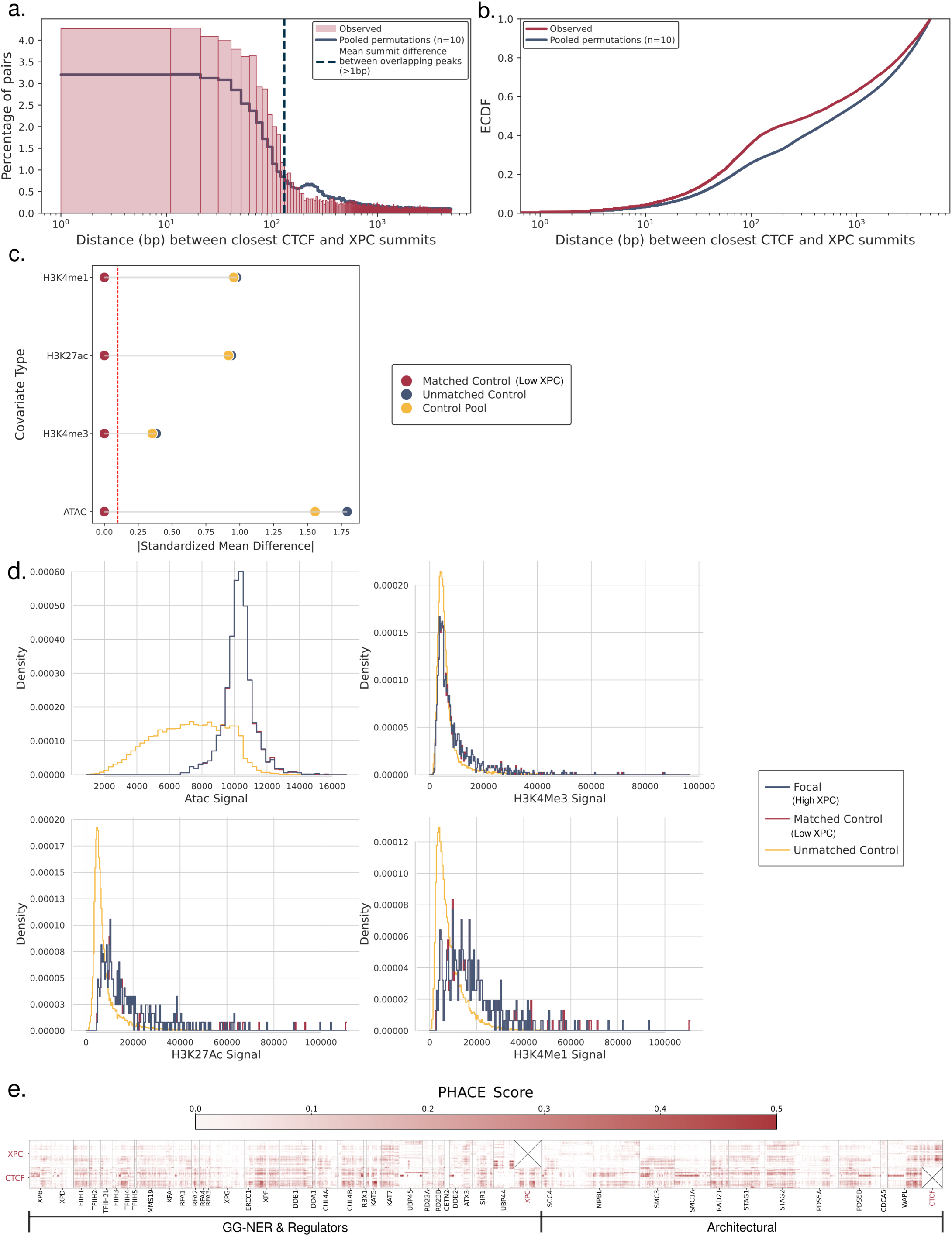
Spatial proximity, covariate-matched validation, and evolutionary coupling between XPC and CTCF. **a,** Distribution of distances from XPC chromatin immunoprecipitation sequencing (ChIP-seq) peak summits to the nearest CTCF ChIP-seq genome-wide peak summits in wild-type (WT) U2OS cells under untreated conditions. The observed distances (red hatched histogram) were compared to a null distribution (blue) generated by repeated permutations of XPC peak positions restricted to ATAC-seq-defined accessible chromatin while preserving chromatin identity, providing an accessibility-matched expectation for XPC-CTCF proximity. **b,** Empirical cumulative distribution function (ECDF) of observed (red) and permuted (blue) distances between CTCF and XPC peak summits. **c,** Covariate balancing diagnostics to match high-XPC to low-XPC bins. Standardized mean differences between matched, unmatched, and control bins are shown for chromatin covariates including histone modifications (H3K27ac, H3K4me1/me3) and chromatin accessibility (ATAC-seq). **d,** Density distributions of chromatin covariates used for matching. **e,** PHACE heatmap summarising residue-level evolutionary coupling between XPC or CTCF and a panel of nucleotide excision repair and chromatin architectural proteins. Each pixel represents a pair of amino-acid positions between the indicated proteins. Scores represent the sum of positive residue-residue coupling signals normalised by the total number of possible amino acid pairings, thereby controlling for protein length. Lack of detectable coupling for certain established interaction partners (e.g. RAD23B with XPC) likely reflects strong evolutionary conservation and limited sequence variation, which reduces the sensitivity of co-evolution-based inference.

**Extended Data Fig. 7.**
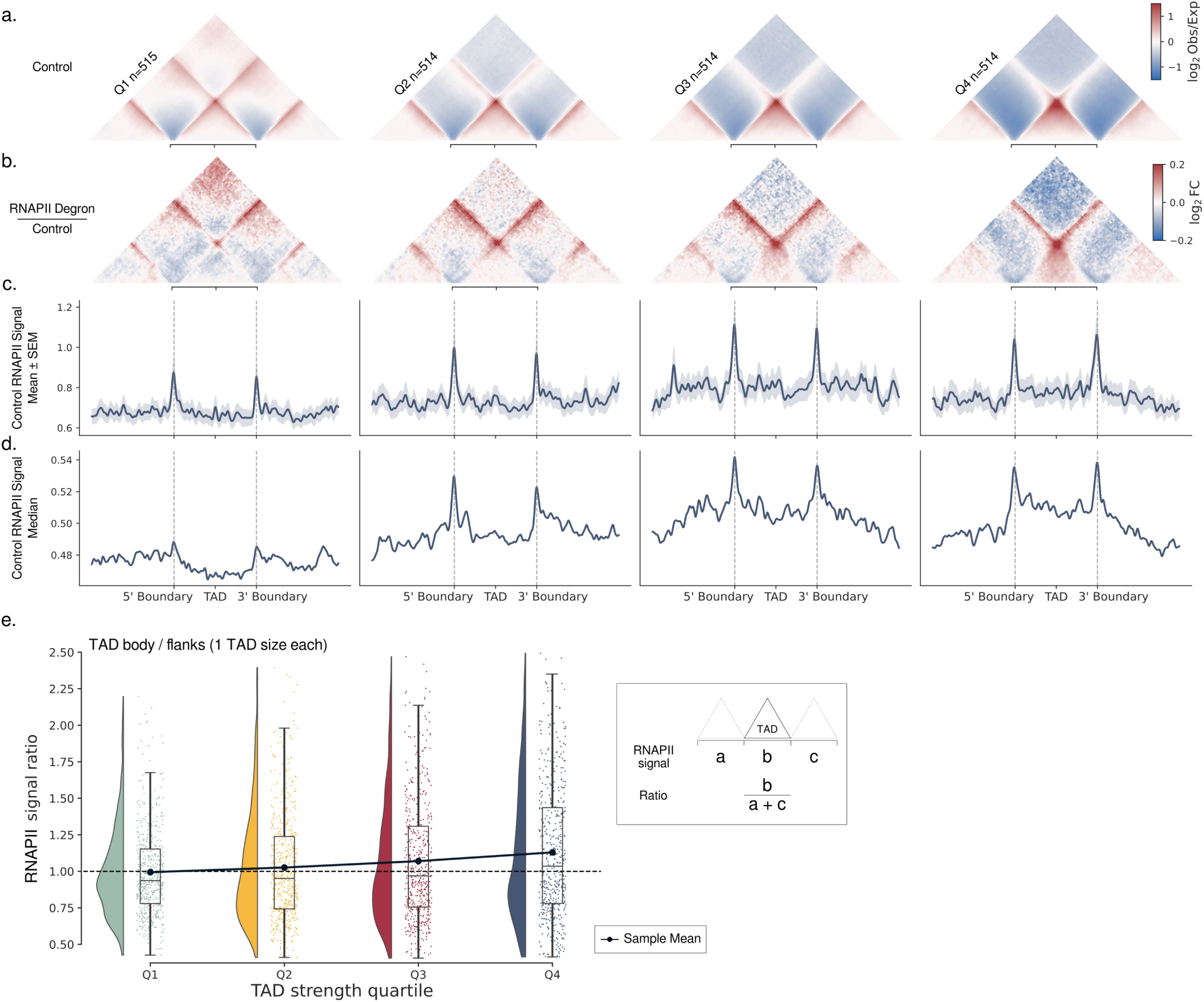
TADs enriched for RNAPII occupancy show increased TAD strength following RNAPII depletion. **a,** TADs, from re-analysed Micro-C data of DLD-1 cells before and after degron-mediated RNAPII depletion^50^, were grouped into quartiles according to TAD strength in the RNAPII-depleted contact map, from weakest (Q1) to strongest (Q4). Aggregated observed-over-expected normalised contacts from the untreated control sample are shown for these TAD groups. **b,** Fold change in normalised contacts after RNAPII depletion relative to the untreated control, shown across the same TAD-strength quartiles. c,d, Untreated RNAPII ChIP-seq signal (from^60^) projected across the same TAD quartiles. Line plots show mean (c) or median (d) signal across regions; shaded bands indicate mean ± SEM. **e,** RNAPII enrichment within the TAD body relative to the flanking regions. Ratio was calculated as the summed RNAPII signal across the TAD body divided by the summed signal across the two flanks, each spanning one TAD length.

**Extended Data Fig. 8.**
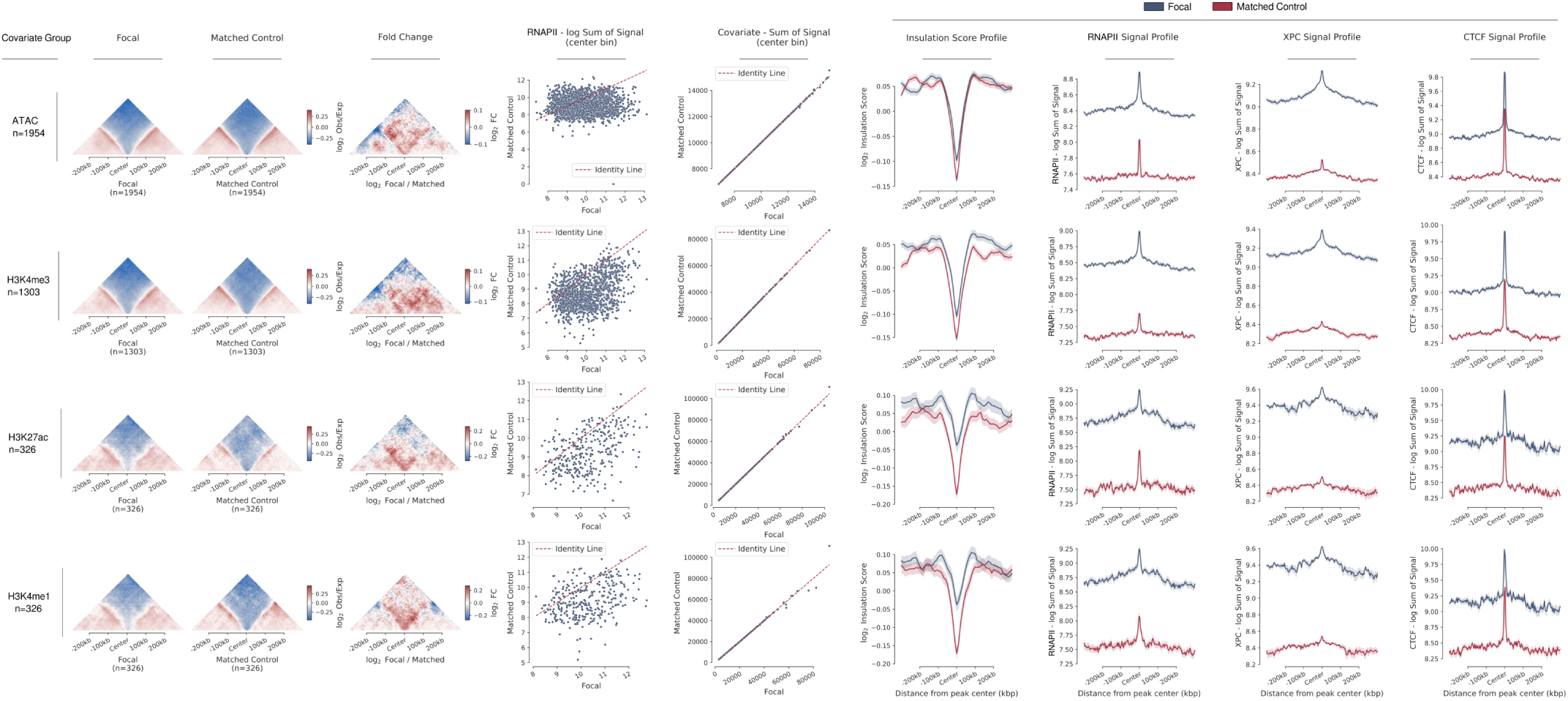
Covariate matching including RNA polymerase II occupancy. For each covariate group, rows show aggregate Hi-C contact maps centred on focal (high-XPC) bins and matched controls (low-XPC) at 10-kb resolution, corresponding log2 fold-change maps (focal-over-matched control), pairwise relationships between RNAPII and the matched covariate, summed covariate signal per bin, and average insulation score, RNAPII, XPC, and CTCF signal profiles across matched regions. Line plots show mean signal across regions; shaded bands, mean ± SEM.

**Extended Data Fig. 9.**
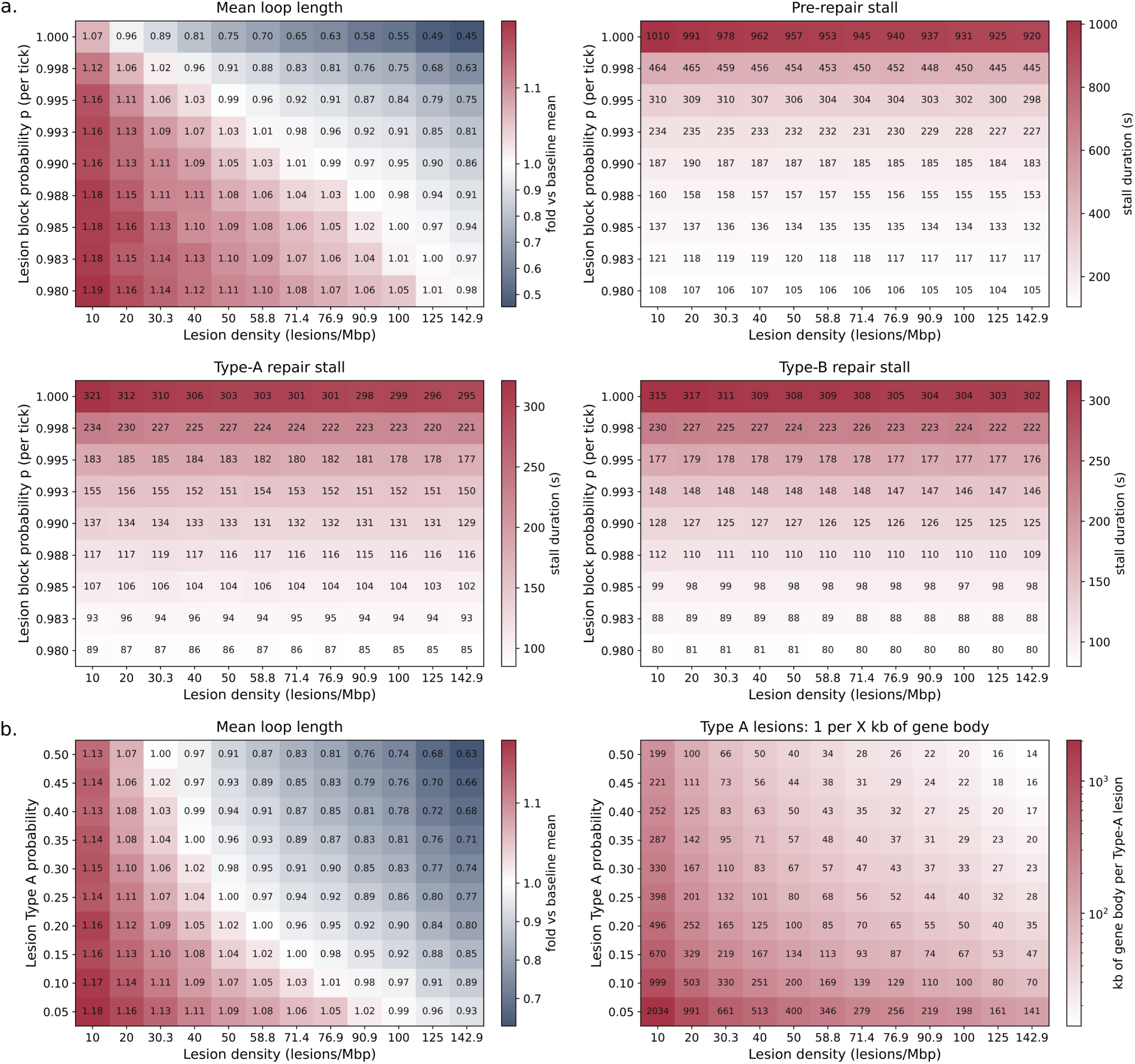
Parameter optimisation for loop extrusion simulations. **a,** Two-dimensional parameter sweep varying lesion density and lesion-associated cohesin stalling probability. Heatmaps show mean loop length (top left), mean duration of Type A pre-repair stalls (top right), mean duration of Type A repair-associated stalls (bottom left), and mean duration of Type B repair-associated stalls (bottom right), measured from simulation trajectories. **b,** Two-dimensional parameter sweep varying lesion density and Type A lesion probability. Heatmaps show mean loop length (left) and the effective frequency of Type A lesions within gene bodies (right), measured from simulation trajectories.

**Extended Data Fig. 10.**
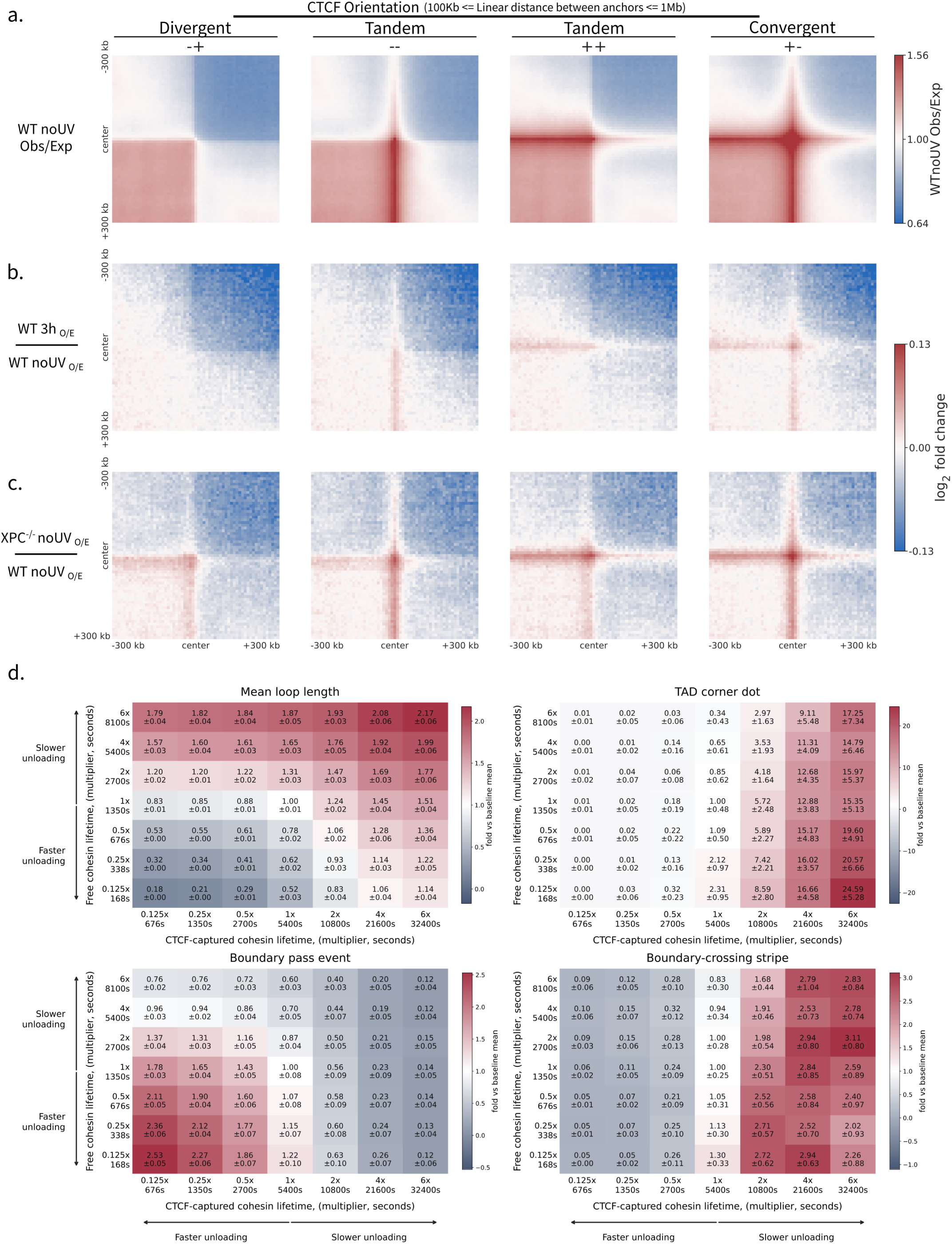
Architectural features of XPC-deficient chromatin and corresponding simulation parameter sweeps. **a,** Aggregate contact maps centred on pairs of CTCF sites identified from IDR-filtered CTCF ChIP-seq peaks shared between undamaged WT cells, WT cells 3 h after UV irradiation, and XPC-deficient (*XPC*^-/-^) cells. Sites were assigned orientation using the nearest CTCF motif occurrence and grouped into divergent (-+), tandem (–), tandem (++), and, as a proxy for TAD boundaries, convergent (+-) pairs. Pileups were generated from all CTCF pairs separated by 100 kb–1 Mb on the same chromosomal arm and show observed-over-expected contact enrichment in undamaged WT cells. **b,** Log2 fold-change contact maps comparing WT cells 3 h after UV irradiation relative to undamaged WT cells. **c,** Log2 fold-change contact maps comparing *XPC*^-/-^ with WT cells. **d,** Two-dimensional parameter sweep varying the residence time of freely extruding cohesin (y-axis) and CTCF-captured cohesin (x-axis). Heatmaps show mean loop length (top left), TAD corner-dot intensity (top right), boundary-pass frequency (bottom left), and boundary-crossing stripe intensity (bottom right), measured from simulation trajectories. Values are shown relative to the baseline simulation.

