## Supplementary Methods for "RNAPII and XPC remodel the 3D genome for UV repair"

One-dimensional stochastic modelling of loop extrusion, transcription and nucleotide-excision repair

**Topics covered**

#### 1 Model overview, time steps and lattice setup

We modelled interphase chromatin as a one-dimensional lattice and simulated the stochastic dynamics of three factors: cohesin loop-extrusion complexes, RNA polymerase II (RNAPII), and ultraviolet photolesions. Building on established stochastic lattice models of loop extrusion (Fudenberg et al., 2016; Banigan et al., 2020) and the polychrom simulation framework (Imakaev et al., 2019), we advanced the system in discrete, fixed-length time steps. Within each step, we updated lesion states, then (when transcription was active) recruited and translocated polymerases, and finally translocated cohesin, under hard occupancy exclusion so that no two factors occupied the same lattice site. State transitions (binding, dissociation, stage changes, capture and release) were memoryless, each drawn as a per-step Bernoulli trial equivalent to first-order (Poisson) processes. We report each transition either as a rate constant or as its characteristic time. We did not simulate the three-dimensional polymer. The features to which population contact maps and single-locus imaging are most sensitive (the one-dimensional cohesin distribution, the lengths of the loops it extrudes, and the sites at which extrusion is impeded) are themselves one-dimensional and can therefore be sampled with far greater statistical depth in this framework. The simulation engine is available at `https://github.com/CompGenomeLab/ner_rnapii_3d`.

Two identities fixed the physical scale. Each lattice site corresponded to 250 bp of DNA, and each time step corresponded to 2 s of real time. All velocities, lifetimes and rates are reported in physical units after applying these identities. Each memoryless transition fired with a per-step probability set by its mean waiting time $\tau$,

$$p \;=\; 1 - e^{-\Delta t/\tau} \;\approx\; \frac{\Delta t}{\tau}, \qquad \Delta t = 2\ \mathrm{s}, \tag{1}$$

Accordingly, each process is specified by a physical rate $1/\tau$ or duration $\tau$ and converted to a per-step probability through Eq. 1, so the modelled kinetics are independent of the chosen step size. The calibrated quantities are reported in Table 1.

We simulated a 10-Mb locus (40,000 sites) instantiated as twenty independent replicate copies (200 Mb of lattice in total), equilibrated each replicate for ≈22 h of simulated time (40,000 steps), and then sampled an ≈89 h production trajectory (160,000 steps). The configurations and analyses are described in Section 8.

**Table 1. Calibrated model parameters.** Values are reported in physical units after applying the lattice (1 site = 250 bp) and temporal (1 step = 2 s) calibrations.

| Parameter | Calibrated value | Basis |
|---|---|---|
| *System* | | |
| Spatial resolution | 250 bp per site | calibration |
| Temporal resolution | 2 s per step | calibration |
| Simulated locus | 10 Mb × 20 replicates (200 Mb) | – |
| Equilibration / production | ≈22 h / ≈89 h | – |
| *Loop extrusion* | | |
| Cohesin separation (density) | one complex per 240 kb | Gabriele et al., 2022 |
| Subunit velocity | 125 $\mathrm{bp\,s^{-1}}$ (7.5 $\mathrm{kb\,min^{-1}}$) | see §3 |
| Loop-extrusion rate | 250 $\mathrm{bp\,s^{-1}}$ (15 $\mathrm{kb\,min^{-1}}$) | ≈2× the in-vivo rate inferred for Gabriele et al., 2022 (≈150 kb / ≈20 min; §3); below in-vitro ≈0.5–2 $\mathrm{kb\,s^{-1}}$ (Kim et al., 2019; Davidson et al., 2019) |
| Residence, free | ≈22.5 min | ≈22 min residence (Hansen et al., 2017); ≈20 min base lifetime (Gabriele et al., 2022) |
| Residence, CTCF-captured | ≈90 min (≈4×) | Gabriele et al., 2022 (stabilised fourfold) |
| Processivity (emergent) | a few $\times 10^2$ kb | cf. ≈150 kb (Gabriele et al., 2022); 300 kb (Yang & Hansen, 2024) |
| Recruitment | uniform | Banigan et al., 2023 |
| *Transcription* | | |
| Promoter-proximal pause | ≈250 bp downstream | – |

| Parameter | Calibrated value | Basis |
|---|---|---|
| Single-Pol pause (median / active) | ≈42 s / ≈25 s | ≈42 s mean live-cell residence (Steurer et al., 2018); ≈1.2–1.4 min model half-life (Zhao et al., 2023); multi-minute by decay-rate (Jonkers et al., 2014; Core & Adelman, 2019) |
| Productive fraction (median; activity-graded) | ≈0.29 (range ≈0.14–0.47) | ≈0.2 (≈80% premature termination), STL-seq (Zimmer et al., 2021) |
| Initiation / loading, expressed (realised) | median ≈0.27 $\text{min}^{-1}$ (occupied genes); ≈0.05 all-gene | low ≈0.2 / high ≈1.0 $\text{min}^{-1}$ (Zhao et al., 2023); pause-initiation limit (Gressel et al., 2019) |
| Productive output, 3′ (realised) | detected median ≈0.02 $\text{min}^{-1}$/allele; active ≈0.1 | ≈2 mRNA $\text{h}^{-1}$ per gene (Schwanhäusser et al., 2011); ≈87 $\text{min}^{-1}$ for heat-shocked HSPA1A (Gressel et al., 2019) |
| Elongation velocity | 2.1 kb $\text{min}^{-1}$ (0.035 kb $\text{s}^{-1}$); ≈1.9 realised | in-vivo 1.3–4.3 kb $\text{min}^{-1}$ (Guo & Price, 2013); 0.5–4 genome-wide (Jonkers et al., 2014); ≈2 assumed by Zhao et al., 2023 |
| 3′ termination dwell | ≈1.8 min (realised) | derived (Fong et al., 2015; Cortazar et al., 2019) |
| *RNAPII – cohesin* | | |
| Cohesin transit past engaged RNAPII | ≈100 s ($k \approx 0.01$ $\text{s}^{-1}$) | Banigan et al., 2023 |
| Cohesin transit past pre-initiation RNAPII | ≈12 s (≈8× more permeable) | – |
| Subunit : elongation velocity ratio | ≈3.6 (loop ≈7) | Banigan et al., 2023 (≥3–5) |
| *Lesions and repair* | | |
| Lesion density (swept) | 1 per 7–100 kb (20 J $\text{m}^{-2}$; repair time-course, Table 2) | §6 |
| TC-NER (Type A) fraction of gene-body lesions | ≤ 0.5 (ceiling); ≈0.20 (typical) | – |
| Pre-repair stage | ≈40 min | – |
| Active repair stage | ≈6 min | – |
| Per-step block, repair stage | ≈0.993 (bypass ≈0.007 / step) | – |

#### 2 Topology: genes, domains and boundaries

We generated the genomic substrate procedurally to reproduce the marginal statistics of a gene-rich human locus. A smooth latent gene-density profile was sampled along the locus and used to partition it into topologically associating domains (TADs) whose lengths were anticorrelated with local gene density: dense regions were tiled by shorter domains (target median ≈0.75 Mb) and sparse regions by longer ones (target median ≈1.25 Mb), with individual lengths drawn from a heavy-tailed distribution (realized range 0.51–1.87 Mb across the 10 domains), bracketing the sub-megabase domain-size distribution of mammalian genomes (median ≈0.88 Mb in mouse ES cells; Dixon et al., 2012). We scaled domains above the characteristic single-loop reach such that the emergent observable reflected domain-level insulation rather than isolated loop corner-peaks.

Each domain boundary was modelled as a permeable, oriented CTCF barrier with a per-encounter probability of arresting an incident cohesin subunit. Boundaries were oriented convergently (the left boundary arrested left-travelling subunits and the right boundary, right-travelling subunits), recapitulating the inward CTCF-motif polarity required for stable looping. Arrest probabilities were modest (calibrated per-encounter values of ≈0.09–0.15, raised severalfold in the insulation sweeps of Section 8) and were set higher at boundaries flanking shorter, denser, more transcriptionally active domains. Each boundary's strength was an equal-weight blend of two drivers, the transcriptional activity and the gene density of its flanking domains, each taken from the more extreme (more active, denser) of the two flanks, so that boundary strength fell with domain length and active, gene-dense domains were well insulated on both sides. Reflecting boundary conditions were imposed at both ends of the locus.

We placed genes at a locus-average density of ≈12 per Mb (122 genes across the 10-Mb locus), with the per-domain count following the same gene-density latent that set the domain partition (such that shorter, denser domains were more gene-dense), and with lengths drawn from a log-normal distribution (median ≈24 kb) with a tail beyond 100 kb, matching the human gene-length distribution. Placement was non-overlapping and interior genes were assigned transcriptional polarity at random, yielding co-directional, convergent and divergent neighbours in their natural proportions. Genes belonged to four regulatory classes: constitutive housekeeping, highly expressed housekeeping, cell-type-specific (the plurality) and developmental, differing in basal output, in the fraction held silent, and in their dependence on a distal enhancer. Each domain carried a transcriptional propensity set primarily by its inverse length and secondarily by its gene density. Gene-class composition was sampled from this propensity, such that shorter domains were enriched for more active classes and longer domains for more poised or silent ones. This resulted in a compartment-like anticorrelation between domain length and transcriptional output.

We recapitulated the enrichment of housekeeping start sites at TAD boundaries (Dixon et al., 2012) as an emergent feature. Approximately half of boundaries were assigned a single promoter positioned 3–25 kb inside the domain interior, with ≈70% designated as housekeeping promoters. This yielded an approximately twofold enrichment of transcription start sites near boundaries while avoiding the artificial peak that would have arisen from placing promoters directly at boundary positions. Regulated-class genes were assigned one or more enhancers, with promoter–enhancer distances drawn from a log-normal distribution (median ≈30 kb, tail extending beyond 500 kb), consistent with measured enhancer–promoter separation distributions, although a minority of pairs spanned a boundary. Productive transcription of these genes required the enhancer and promoter to be simultaneously contained within a cohesin loop (Section 4), thereby recapitulating the cohesin-dependent nature of enhancer–promoter communication observed for Mediator–cohesin looping

(Kagey et al., 2010) and for enhancer-driven transcriptional activation (Narita et al., 2025).

### 3 Loop extrusion

We modelled cohesin as a loop-extruding complex with two motor subunits that bridge the base of a chromatin loop (the stretch of lattice between the two subunits constituting the loop). A complex initially occupied a pair of adjacent sites and extruded a loop by translocating its two subunits bidirectionally away from one another, each advancing one site per time step (symmetric, two-sided extrusion). As in related cohesin simulations (Gabriele et al., 2022; Banigan et al., 2020), complexes could not bypass one another and could not translocate beyond the terminal lattice sites. A subunit advanced only into a vacant site; a boundary, another cohesin, a polymerase or a lesion arrested that subunit while the contralateral subunit could continue, such that the complex extruded asymmetrically when one side was blocked.

With the 2 s time step, one-site-per-subunit translocation gives a subunit velocity and a loop-extrusion rate of

$$v_{\mathrm{sub}} = \frac{\ell}{\Delta t} = \frac{250\ \mathrm{bp}}{2\ \mathrm{s}} = 125\ \mathrm{bp\,s^{-1}}, \qquad v_{\mathrm{loop}} = 2\,v_{\mathrm{sub}} = 250\ \mathrm{bp\,s^{-1}}. \tag{2}$$

This loop-extrusion rate is about twofold faster than the ≈125 $\mathrm{bp\,s^{-1}}$ that the in-vivo measurements imply for the loop as a whole (the ratio of the ≈150 kb processivity to the ≈20 min residence reported by Gabriele et al., 2022; the paper does not state an extrusion speed directly), yet below the ≈0.5–2 $\mathrm{kb\,s^{-1}}$ measured in single-molecule assays (Kim et al., 2019, mean ≈0.5 $\mathrm{kb\,s^{-1}}$; Davidson et al., 2019, up to ≈2.1 $\mathrm{kb\,s^{-1}}$); it therefore lies between the in-vivo and in-vitro estimates. (Equivalently, our 2 s step is half the ≈4 s that the in-vivo rate would imply.) We adopted the faster rate such that each extruding subunit outpaced transcription elongation (2.1 $\mathrm{kb\,min^{-1}}$; Section 4) by ≈3.6-fold (and the loop as a whole by ≈7-fold), at or above the at-least-threefold-to-fivefold separation required to reproduce transcription's effect on contact maps (Banigan et al., 2023).

We held the cohesin complement at a fixed mean density of one complex per 240 kb (≈833 complexes across the twenty replicates), as estimated for mammalian cells (Gabriele et al., 2022). Each complex unbound stochastically with a per-step probability set by its mean residence time (Eq. 1), and a complex that dissociated immediately reloaded at a random unoccupied position, holding the total constant. The free residence time was ≈22.5 min, matching the ≈22 min cohesin residence measured by live-cell imaging (Hansen et al., 2017) and near the ≈20 min base lifetime adopted by Gabriele et al., 2022; the product of extrusion rate and residence gives an emergent (unobstructed) processivity of a few hundred kilobases, on the scale of the ≈150–300 kb adopted in recent extrusion models (≈150 kb inferred in vivo by Gabriele et al., 2022; 300 kb used by Yang & Hansen, 2024):

$$N_{\mathrm{LEF}} = \frac{L_{\mathrm{tot}}}{d} = \frac{200\ \mathrm{Mb}}{240\ \mathrm{kb}} \approx 833, \qquad \lambda \approx v_{\mathrm{loop}}\,\tau_{\mathrm{res}} \approx 15\ \mathrm{kb\,min^{-1}} \times 22.5\ \mathrm{min} \approx 340\ \mathrm{kb}, \tag{3}$$

with $L_{\mathrm{tot}}$ the total lattice length, $d$ the cohesin separation and $\tau_{\mathrm{res}}$ the free residence time. This product is an unobstructed upper bound; as boundaries, collisions and polymerase encounters truncate loops, the realised mean loop length in the transcription-on simulations was ≈216 kb, within the ≈150–300 kb range. We recruited complexes uniformly at random, without preference for promoters or enhancers: uniform loading reproduces contact maps better than promoter-targeted loading, and cohesin enrichment at active genes arises downstream from polymerase barriers rather than from biased recruitment (Banigan et al., 2023; Busslinger et al., 2017).

A subunit reaching an oriented boundary was captured (arresting extrusion on that side) with the boundary's per-encounter probability, and otherwise passed through; boundaries therefore biased rather than absolutely blocked extrusion. A captured complex was stabilised against turnover, extending its residence ≈4-fold, to ≈90 min (four times the ≈22.5 min free residence), matching the ≈4-fold CTCF-dependent stabilisation of cohesin reported in vivo (Gabriele et al., 2022), and reflecting the distinct chromatin-residence dynamics of CTCF and cohesin (Hansen et al., 2017).

### 4 Transcription

We represented transcription at single-molecule resolution as the life cycle of individual RNAPII complexes, because the translocating polymerase, not a scalar expression level, interacts sterically with cohesin. A polymerase was recruited stochastically at a promoter and entered an obligatory promoter-proximal pause one lattice site (≈250 bp) downstream of the start site (the single-site lattice resolution, coarser than the physiological ∼30–60 bp pause position). A paused polymerase *occluded* the promoter: while it occupied the promoter-proximal window, it suppressed the loading of a new polymerase; thus gene throughput was limited by the rate of pause escape rather than by recruitment, the pause–initiation limit (Gressel et al., 2019). We therefore controlled output through pause-escape kinetics; increasing the recruitment rate alone only increased promoter-proximal occupancy without raising productive output.

A paused polymerase resolved in one of two competing ways: it either escaped into productive elongation or terminated prematurely and was lost. The probability of productive escape (the productive fraction) was specified on a per-gene basis set near the ≈20% inferred for mammalian promoters, where roughly 80% of promoter-proximally paused RNAPII terminate prematurely (Zimmer et al., 2021, by STL-seq in human and *Drosophila* cells; consistent with the rapid turnover of promoter-paused RNAPII seen by live imaging, Steurer et al., 2018). To reflect the lower premature-termination rates of highly expressed genes, we made the productive fraction increase with transcription activity, ranging from ≈0.14 for weakly transcribed genes to ≈0.47 for the most active genes, with a population median of ≈0.29.

A single parameter determines how long a paused polymerase remains at the promoter before either escaping into productive elongation or terminating. We set the pause dwell time to ≈42 s, matching the promoter-proximal residence time of endogenous RNAPII measured by live-cell imaging in human cells (Steurer et al., 2018). To reflect differences in transcriptional activity, dwell times were scaled inversely with gene expression, such that the most active genes cleared the pause within ≈25 s whereas poised genes remained paused for longer. Population-based measurements have reported much longer pause durations, including multi-minute half-lives inferred from transcription-inhibitor decay assays (mean ≈7 min in mouse ES cells, Jonkers et al., 2014; ≈2–30 min across studies, Core & Adelman, 2019). These estimates exceed single-molecule dwell times because population-level pause duration depends strongly on the metric used. In the model, although individual polymerases dwell for only ≈42 s, dividing pause occupancy by productive output, which is reduced by the large fraction of polymerases that terminate before completing transcription, yields an apparent pause duration of ≈2.6 min. A single-second scale of dwell time is therefore consistent with both single-molecule measurements and longer population-level estimates.

We drew per-gene activity from a heavy-tailed, power-law distribution spanning two to three orders of magnitude, approximating the broad, continuous distribution of transcription rates observed across genes. Each gene's polymerase-loading propensity was determined by three factors: its pause-escape probability, a weak inverse power dependence on domain length (such that genes

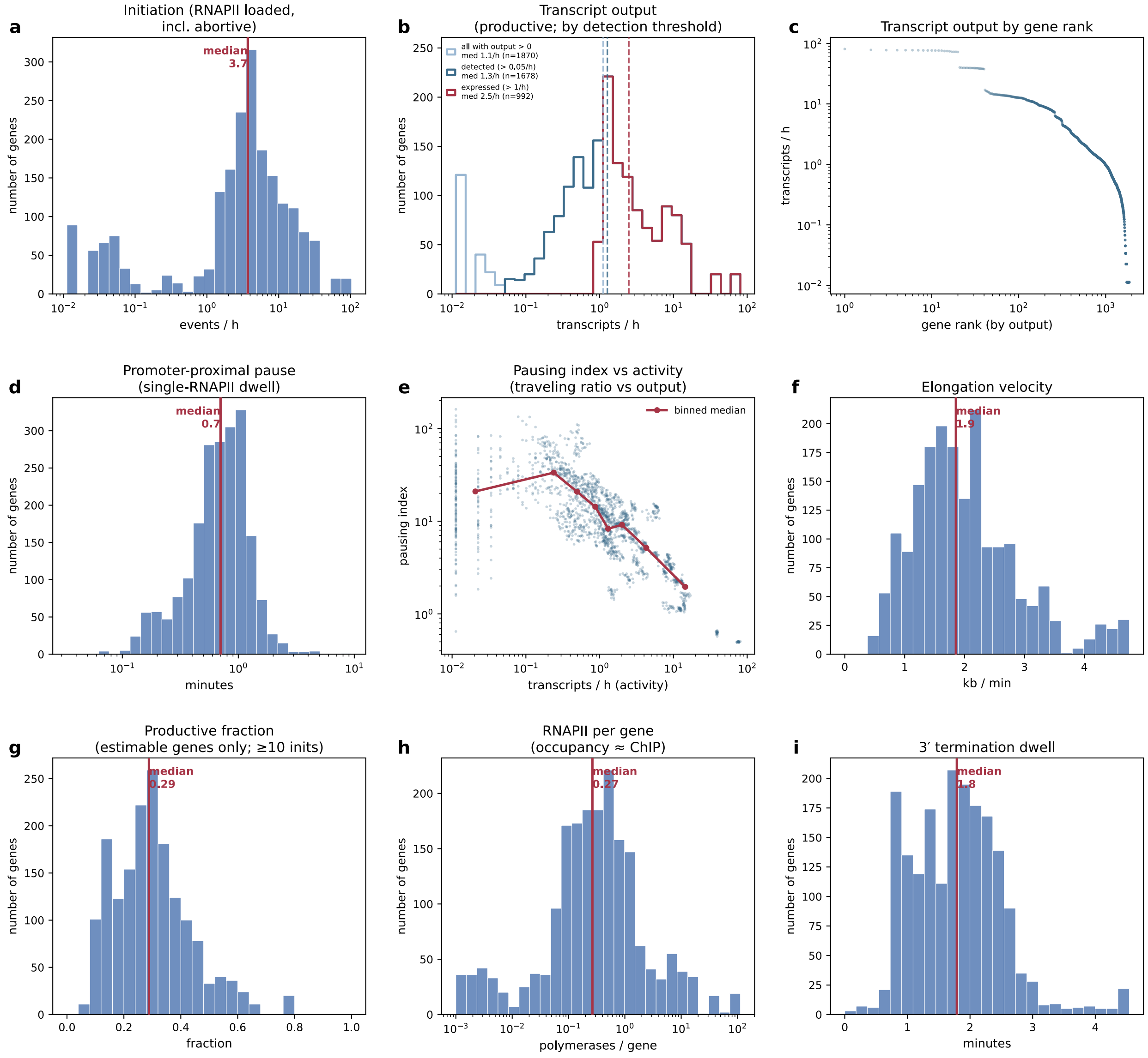

Figure 1: **Per-allele transcription kinetics across the simulated allele population.** Distributions were calculated from 2,440 allele-instances (122 loci (genes) across 20 replicate chains). Red lines indicate model medians. **(a)** Initiation rate, defined as RNAPII loading events, including the 70–80% of polymerases that terminate prematurely during promoter-proximal pausing. **(b)** Productive transcript output, defined as RNAPII molecules reaching the 3′ ends of genes. Nested step histograms are shown for several detection thresholds (output $>0$, $>0.05$ h$^{-1}$, $>1$ h$^{-1}$), each with its own dashed median, such that the "expressed median" is cutoff-dependent. **(c)** Productive transcript output by gene rank (log–log scale). **(d)** Promoter-proximal pause dwell time of individual polymerases. **(e)** Pausing index versus productive transcript output (log–log scale). The red line shows the binned-median trend. Highly expressed genes show lower pausing indices. **(f)** Elongation velocity. **(g)** Productive fraction, shown only for genes with at least 10 initiation events, for which the ratio can be estimated reliably. Genes with fewer than 10 initiations (low-confidence) or no initiations (undefined ratio) are tallied separately and excluded from the distribution. **(h)** RNAPII occupancy per gene. **(i)** 3′ termination dwell time. Heavy-tailed distributions are displayed on logarithmic axes.

in shorter, more active domains recruited polymerase more often) and a power-law-distributed stochastic scatter term, renormalised such that the geometric-mean loading of the expressed set met a fixed target and was subsequently scaled by an absolute per-class factor:

$$r_i = r_0 \frac{\rho_i}{\left(\prod_{j\in\mathcal{E}} \rho_j\right)^{1/|\mathcal{E}|}} c_{c(i)}, \qquad \rho_i = p_i^{\mathrm{esc}} \left(\frac{\tilde{L}}{L_{T(i)}}\right)^{\eta} \pi_i, \quad \pi_i \sim \mathrm{PowerLaw}(\gamma), \tag{4}$$

where $r_i$ is gene $i$'s loading propensity, $r_0$ the target geometric-mean loading over the expressed set $\mathcal{E}$, $c_{c(i)}$ its regulatory-class multiplier, $p_i^{\mathrm{esc}}$ its pause-escape probability, $L_{T(i)}$ its domain length, $\tilde{L}$ the median gene-hosting domain length, $\eta \approx 0.2$ a weak size exponent and $\pi_i$ a power-law (Zipf) scatter term (exponent $\gamma \approx 2.4$). A defined fraction of the regulated-class genes was held effectively silent. Rather than assigning these genes a loading rate of zero, we drew their loading propensities from a graded, near-zero range extending from deeply silent (no initiation over the trajectory) to a low basal ("leaky") level. This produced a continuous transition between silent and expressed genes rather than an artificial spike-and-gap, consistent with real transcriptomes.

Escaped polymerases translocated in their transcriptional direction at a gene-specific velocity drawn around a 2.1 $\mathrm{kb\,min^{-1}}$ anchor (0.035 $\mathrm{kb\,s^{-1}}$; per-gene range ≈0.5–6 $\mathrm{kb\,min^{-1}}$), within the 1.3–4.3 $\mathrm{kb\,min^{-1}}$ reported for mammalian RNAPII in vivo (Guo & Price, 2013) and bracketing the 0.5–4 $\mathrm{kb\,min^{-1}}$ measured genome-wide in mouse ES cells (Jonkers et al., 2014). Steric exclusion among co-transcribing polymerases reduced the realised median to ≈1.9 $\mathrm{kb\,min^{-1}}$, close to the rate assumed in genome-wide initiation inference (Zhao et al., 2023). Multiple polymerases could co-occupy a gene subject to steric exclusion, such that inter-polymerase spacing emerged self-consistently. On reaching the $3'$ end, a polymerase decelerated and dwelt over the termination zone for ≈1.8 min before release. This dwell represents the multi-kilobase termination window downstream of the poly(A) site (Fong et al., 2015), across which RNAPII decelerates from >2 to <1 $\mathrm{kb\,min^{-1}}$ (Cortazar et al., 2019), and captures the cohesin accumulation associated with terminating polymerases at $3'$ gene ends (Banigan et al., 2023). For regulated-class genes, pause escape was permitted only while the gene's promoter and a cognate enhancer were co-contained within a single cohesin loop, gating a subset of productive initiation on the instantaneous loop-extrusion state and reproducing the cohesin-dependence of enhancer-driven activation established for Mediator–cohesin looping (Kagey et al., 2010) and enhancer–promoter communication (Narita et al., 2025).

Across the population (122 genes × 20 replicate chains = 2,440 allele-instances; Fig. 1), these kinetics reproduced genome-scale transcription statistics. About 69% of alleles were detectably transcribed (productive output >0.05 $\mathrm{transcripts\,h^{-1}}$); the rest formed a continuum from deep-silent to low basal output, with ≈23% producing no completed transcript and ≈17% never initiating at all. The typical detected allele made ≈1.3 $\mathrm{transcripts\,h^{-1}}$ (≈2.6 per diploid gene), close to the ≈2 $\mathrm{mRNA\,h^{-1}}$ average synthesis rate measured genome-wide in mammalian cells (Schwanhäusser et al., 2011). Per-gene output was heavy-tailed, with a thin active tail reaching tens of $\mathrm{transcripts\,h^{-1}}$. Promoter-proximal pausing was prevalent, with a median pausing (travelling-ratio) index of ≈6 over expressed genes, in line with the widespread pausing of mammalian genes (Day et al., 2016). The modelled locus is gene-rich and transcriptionally active, with expression levels above the genome-wide median. However, its global output remains well below the *hypertranscription* of cancer cells, where total RNA production is increased approximately twofold on average across human cancers and by as much as 3.5-fold in aggressive subtypes (Zatzman et al., 2022). These values describe genome-wide amplification of RNA output rather than per-allele transcription rates. Such hypertranscription is frequently associated with elevated MYC activity (Lin et al., 2012), although MYC regulates transcription in a substantially gene-selective manner rather than acting as a universal amplifier (Sabò et al., 2014; Walz et al., 2014). The most active genes in simulation accordingly fall short of cancer-line single-gene maxima: our top alleles reach ≈1.3 productive events $\mathrm{min}^{-1}$ *per allele* (≈2.6 per diploid gene), whereas a strongly induced gene such as HSPA1A reaches ≈87 events $\mathrm{min}^{-1}$ *per cell* upon heat-shock induction in the hyper-transcribing human line K562 (Gressel et al., 2019). Model transcription rates are reported on a per-allele basis throughout. Our parameters thus describe an active, non-malignant locus, with the overall output level controlled by a single loading-rate parameter.

### 5 RNAPII – cohesin interactions

We treated an engaged polymerase as a permeable, motile barrier to extrusion (Banigan et al., 2023), resolving two encounter geometries. When a cohesin subunit attempted to step onto a polymerase-occupied site, it succeeded only stochastically. The mean transit time past a transcriptionally engaged (paused, elongating or terminating) polymerase was ≈100 s, corresponding to a bypass rate of ≈0.01 $\mathrm{s}^{-1}$, which best reproduces the effects of transcription on contact maps (Banigan et al., 2023). Two quantities define this barrier, the per-step probability that a subunit is blocked rather than passing, and the velocity margin that makes the barrier directional:

$$p_{\mathrm{block}} = e^{-\Delta t/\tau_{\mathrm{bypass}}}, \qquad \tau_{\mathrm{bypass}}^{\mathrm{engaged}} \approx 100 \text{ s } (p_{\mathrm{block}} \approx 0.98), \qquad \frac{v_{\mathrm{sub}}}{v_{\mathrm{elong}}} \approx 3.6. \tag{5}$$

A merely pre-initiation polymerase was ≈8-fold more permeable ($\tau_{\mathrm{bypass}} \approx 12$ s). We did not accelerate the turnover of a cohesin held against a polymerase: its unbinding rate remained the baseline value. The framework permits an accelerated cohesin-eviction regime, analogous to the WAPL-mediated release that removes cohesin from active genes in vivo (Busslinger et al., 2017). However, short eviction lifetimes depleted cohesin from active genes rather than generating the experimentally observed enrichment. We therefore did not employ the accelerated eviction regime. Gene-body and 3′ enrichment instead arose entirely from the moving-barrier mechanism described below.

Reciprocally, an elongating polymerase that collided with a cohesin subunit most often relocated it (displacing the subunit ahead and translating the loop along the gene) when the two converged head-on, and more often arrested itself when co-directional; non-elongating polymerases did not relocate cohesin. This rectified interaction drove cohesin accumulation at and beyond 3′ ends and the formation of cohesin islands between convergent genes (Banigan et al., 2023; Busslinger et al., 2017). In the polymerase-free configurations (the transcription-off control and the repair experiments), these interactions were absent and cohesin responded only to boundaries, other cohesins, and lesions.

### 6 Lesions

We represented ultraviolet photoproducts, predominantly cyclobutane pyrimidine dimers and 6-4 photoproducts, as damaged lattice sites. These lesions, the principal substrates of nucleotide-excision repair (NER), were induced at a controlled density and were made to undergo a two-stage repair process before excision. We maintained the population homeostatically at a fixed target $M$ equal to the lattice length divided by the chosen lesion spacing $s$ (the experimental analogue of UV irradiation; below),

$$M = \left\lfloor \frac{L_{\text{tot}}}{s} \right\rfloor. \tag{6}$$

At each simulation step, we progressed existing lesions through their repair states, removed lesions that had completed repair, and spawned new lesions until the target abundance was restored, maintaining a constant steady-state damage burden despite continual lesion turnover. At the start of each simulation, the lesion population was seeded directly by the spawner.

We spawned each new lesion by weighted random sampling of lattice sites. Every site $i$ carried a weight equal to the length of its host domain raised to a negative tunable power, and its induction probability was that weight normalised over the lattice,

$$P(\text{induce at site } i) = \frac{L_{T(i)}^{-\alpha}}{\sum_j L_{T(j)}^{-\alpha}}, \qquad \alpha = 1, \tag{7}$$

where $L_{T(i)}$ is the length of the domain containing site $i$ and $\alpha$ the steepness of the size bias (the configuration key `lesion_tad_size_exponent`; here $\alpha = 1$). The per-tick deficit was drawn in batches and rejection-sampled (sites already carrying a lesion were rejected and redrawn). Because the weights are normalised, any reference length cancels, leaving only relative domain lengths. The exponent sets how the fixed damage budget $M$ partitions across domains: a domain of length $L$ spans $L$ sites, each weighted $L^{-\alpha}$, so its expected share of lesions scales as $L \cdot L^{-\alpha} = L^{1-\alpha}$. At $\alpha = 0$, induction is spatially uniform, with a domain's lesion count growing in proportion to its length, at constant per-kilobase density. At $\alpha = 1$, the value we used, every domain accrues in expectation a similar *number* of lesions regardless of size, such that the per-kilobase damage density scales as $1/L$ and shorter domains are proportionally more heavily damaged. At $\alpha > 1$, shorter domains would carry more damage even in absolute count. All newly introduced lesions entered the pre-repair state. The same inverse-domain-length bias, through the exponent $\beta$ of Eq. 10, also accelerates recognition and repair in shorter domains; thus chromatin context shapes both ends of the lesion life cycle.

Lesion spacing is the experimental analogue of UV irradiation; our wet-lab experiments used 20 $\mathrm{J\,m^{-2}}$ UVC, which generates ≈1 photolesion (CPD and 6-4PP combined) per 15 kb of single-stranded DNA (Quinet et al., 2016, after Andrade-Lima et al., 2015), i.e. ≈1 per 7.5 kb of duplex DNA (derived from the per-strand value). With the assumption that UV photoproduct yield is approximately linear in dose over this range, the equivalent dose of any spacing follows from this anchor by inverse proportionality:

$$D \approx D_0 \frac{s_0}{s} \approx \frac{150 \text{ kb} \cdot \mathrm{J\,m^{-2}}}{s}, \qquad (D_0,\, s_0) = (20 \ \mathrm{J\,m^{-2}},\ 7.5 \text{ kb}). \tag{8}$$

with $D$ the equivalent UVC dose, $s$ the lesion spacing (in kb) and $(D_0,\, s_0)$ the 20 $\mathrm{J\,m^{-2}}$ anchor. After irradiation, nucleotide-excision repair clears lesions over hours to days, lowering their genomic density. We modelled this time-course as a sweep of the spacing from ≈7 kb (the fresh ≈22 $\mathrm{J\,m^{-2}}$ burden) up to ≈100 kb (largely repaired), running each spacing as an independent steady-state simulation, a snapshot of the lesion burden at one post-exposure time (Table 2).

**Table 2. Lesion-density calibration and repair-time sweep.** Top row: duplex-DNA photoproduct yield at the 20 $\mathrm{J\,m^{-2}}$ experimental dose (the initial burden). Bottom row: the swept lesion-spacing range, modelling the post-exposure repair time-course; Eq. 8 gives the equivalent dose of each density.

| Lesion spacing (duplex DNA) | Lesion density | UVC dose | Basis |
|---|---|---|---|
| 1 per ≈7.5 kb | 133 lesions $\mathrm{Mb}^{-1}$ | 20 $\mathrm{J\,m^{-2}}$ | Quinet et al., 2016 (1 per 15 kb / strand); Andrade-Lima et al., 2015 |
| 1 per 7–100 kb (swept) | 143–10 lesions $\mathrm{Mb}^{-1}$ | | this study (repair time-course via decreasing densities over independent simulations) |

We assigned each lesion to a repair pathway at the moment of induction. A lesion within a gene body was classified as a transcription-coupled repair lesion (TC-NER; "Type A") with a fixed probability (the TC-NER fraction) and otherwise as a global-genome repair lesion (GG-NER; "Type B"); lesions outside gene bodies were always assigned to GG-NER,

$$P(\text{TC-NER} \mid i) = \begin{cases} f_A, & i \in \text{gene body}, \\ 0, & \text{otherwise}, \end{cases} \qquad f_A \leq \tfrac{1}{2}, \tag{9}$$

with the complement assigned to GG-NER. TC-NER lesions represent damage on the transcribed strand of an active gene, where an elongating polymerase arrests. The stalled polymerase both designates the lesion for rapid, dedicated repair and acts as the extrusion barrier represented in the model. GG-NER lesions comprise intergenic damage and the non-transcribed strand or non-stalling remainder within genes and are recognised by genome-wide surveillance. We treated the TC-NER fraction as a swept parameter. Because only lesions on the transcribed strand of an active gene generate polymerase-stalling events, its theoretical upper bound is 0.5, whereas a value near 0.20 is more biologically realistic owing to the predominance of slowly excised cyclobutane dimers. In control simulations lacking global-genome repair, the expected number of TC-NER lesions was held constant and lesions were sampled exclusively from gene bodies, ensuring that the TC-NER burden matched that of the full-repair simulations rather than being artificially inflated.

Each lesion then followed a two-state, memoryless repair trajectory with no internal countdown, analogous to the treatment of cohesin turnover. At every step, pre-repair lesions advanced to the repair state and repair-state lesions were excised and removed, with per-step probabilities

$$p_{\text{PRE}\to\text{REP}} = \min\Big(1,\ \frac{\Delta t}{\tau_{\text{pre}}}\,\mu_i\Big), \qquad p_{\text{REP}\to\varnothing} = \min\Big(1,\ \frac{\Delta t}{\tau_{\text{rep}}}\,\mu_i\Big), \qquad \mu_i = \Big(\frac{\bar{L}}{L_{T(i)}}\Big)^{\beta}, \tag{10}$$

with mean pre-repair time $\tau_{\text{pre}} \approx 40$ min, mean repair time $\tau_{\text{rep}} \approx 6$ min, $\bar{L}$ the mean domain length and $\beta = 1$. The multiplier $\mu_i$ exceeds one in shorter-than-average domains and is less than one in longer domains (the per-step probability itself being capped at one, via the min). Consequently, lesions in shorter domains were recognised and repaired more rapidly, whereas those in longer domains were repaired more slowly. Thus, the same chromatin-context dependence governed both lesion induction and lesion repair. The stage durations and the lesion density jointly set the steady-state probability that any site carries an obstructing lesion.

#### 7 Lesion – cohesin interactions

In the lesion simulations, transcription was disabled and no transcribing polymerases were explicitly modelled. Instead, a transcription-coupled (Type A) lesion in its pre-repair stage served as a proxy for a polymerase stalled at the damage, providing an equivalent obstacle to cohesin extrusion at the same genomic site. We adopted this representation rather than explicitly modelling polymerase stalling, removal, repair-coupled restart and re-initiation because the kinetics of transcription recovery after UV damage remain insufficiently constrained to support reliable parameterisation. The lesion field therefore represents the steady-state extrusion barrier imposed by stalled polymerases and associated repair complexes without assuming a specific transcription-recovery timescale.

A lesion's effect on cohesin was conditioned on its repair stage and pathway. A lesion in the active repair stage of either pathway arrested an incident cohesin subunit with high per-step probability (≈0.993; bypass ≈0.007 per step), representing the bound repair machinery occluding the duplex. A TC-NER lesion in the pre-repair stage arrested cohesin only in the polymerase-free configurations, where it substituted for the stalled polymerase that would otherwise provide the obstruction. When polymerases were simulated, the block was supplied by the explicitly modelled stalled polymerase (Section 5) and the lesion did not independently arrest cohesin in this stage, avoiding double counting. A GG-NER lesion in the pre-repair stage was transparent to cohesin because no obstructing complex is present prior to assembly of the repair machinery. An arrested subunit remained stalled until either the lesion was removed upon repair completion or the cohesin complex turned over at the baseline rate (no accelerated eviction, as for a polymerase barrier; Section 5). Because only the leading subunit was tested at each step, a cohesin complex continued extruding on its unobstructed side while the opposite subunit remained arrested, allowing lesion fields to reshape rather than abolish the distribution of cohesin loops.

### 8 Simulation pipeline

All results were produced by a single driver, executing the configurations and analyses in a fixed sequence under the calibration of Section 1 (250 bp per site, 2 s per step; ≈22 h equilibration and ≈89 h production per run).

We generated a 10-Mb locus comprising 122 genes distributed across 10 domains and instantiated it as twenty independent replicates. From each replicate we generated two matched configurations: a transcription-on baseline containing active polymerases and a transcription-off control with identical genes, domains and boundaries but no polymerases. Comparison of these configurations isolates the effects of transcription at fixed chromosomal architecture.

Relative to the transcription-off control, transcription reduced the mean loop length (from ≈248 to ≈216 kb), increased cohesin occupancy within gene bodies (≈1.1-fold), and decreased cohesin occupancy at CTCF boundaries (≈1.3-fold) and domain-spanning corner dots (≈4.4-fold; all $P < 0.001$ across 20 replicates), the moving-barrier signature of transcription on chromosome folding (Banigan et al., 2023). Across both conditions we quantified the cohesin-distribution metrics defined in Table 3, including mean loop length, cohesin occupancy at gene bodies and at CTCF boundaries, boundary-pass (read-through) events, boundary-crossing stripes and corner-dot frequency.

To examine the effect of insulation strength on cohesin distribution, we varied boundary arrest probabilities up to ≈6-fold relative to their calibrated values. In polymerase-free simulations, lesions were introduced and three two-dimensional parameter sweeps were performed: TC-NER (polymerase-stalling) fraction versus lesion density, per-step lesion-blocking probability versus lesion density, and recognition and repair timescales versus density. In these experiments the pre-repair stage lasted ≈40 min, the repair stage ≈6 min, the per-step block probability was ≈0.993, and boundary strengths were fixed at 3.25 times their calibrated values.

Finally, keeping transcription active in both the reference and perturbed simulations, we mapped the cohesin-distribution metrics across a two-dimensional grid spanning free cohesin residence time and CTCF-captured (WAPL-protected) residence time. Each parameter was varied from one-eighth to sixfold of its calibrated value, and results were reported as fold changes relative to the transcription-on baseline. This analysis separated the contributions of extrusion processivity and CTCF stabilisation to loop length and boundary occupancy.

**Table 3. Cohesin-distribution metrics.** All metrics are computed from the one-dimensional cohesin trajectories (one complex = two subunits; 1 site = 250 bp; 2 s per sampled frame) and averaged over all sampled frames and the twenty replicates; a site counts as a CTCF anchor when it lies in the oriented capture window of a domain boundary.

| Metric | Definition |
|---|---|
| Mean loop length | The size of the chromatin loop held by a cohesin complex (the genomic distance between its two subunits), averaged over every complex and every sampled frame (kb). It reports the typical extruded loop size. |
| Cohesin at gene bodies | The mean cohesin occupancy of gene bodies: every cohesin subunit lying inside a gene body is counted, then divided by the number of gene-body sites and the number of sampled frames. A high value reflects cohesin enrichment within transcribed genes. |
| Cohesin at CTCF boundaries | The same occupancy measure evaluated at CTCF-anchor sites (mean subunits per anchor site per frame). A high value reflects cohesin accumulation (parking) against convergent CTCF boundaries. |
| Boundary-pass events | How often extrusion reads through a boundary instead of being arrested: for each internal boundary, the rate of single-site subunit steps that cross it, per boundary per time step (genuine one-site moves only; complexes that unbind and reload elsewhere are excluded). It measures boundary permeability. |
| Boundary-crossing stripe | A one-sided loop that straddles a boundary: one subunit stays captured at a CTCF anchor while the other extrudes past a domain boundary, leaving the loop with a single anchored end (counted per boundary per frame). This asymmetric configuration produces an architectural stripe on a contact map; by contrast, a corner dot has both subunits anchored. |
| Corner dot | A loop that pins an entire domain: one subunit captured at the domain's left anchor and the other at its right anchor, bridging it end to end (counted per internal domain per frame). It is the one-dimensional counterpart of the focal corner dot seen at a domain's apex on a contact map. |
